# Cooption and Specialization of Endothelin Signaling Pathways Drove Elaboration of the Neural Crest in Early Vertebrates

**DOI:** 10.1101/710475

**Authors:** Tyler A. Square, David Jandzik, James L. Massey, Marek Romášek, Haley P. Stein, Andrew W. Hansen, Amrita Purkayastha, Maria V. Cattell, Daniel M. Medeiros

**Affiliations:** Department of Ecology and Evolutionary Biology, University of Colorado, Boulder, CO 80309, USA; Department of Molecular and Cellular Biology, University of California, Berkeley, CA 94720, USA; Department of Zoology, Charles University in Prague, Prague, 128 44, Czech Republic; Department of Zoology, Comenius University in Bratislava, Bratislava, 84215, Slovakia

## Abstract

The neural crest (NC) is a vertebrate-specific embryonic tissue that forms an array of clade-defining adult features. A key step in the formation of these diverse derivatives is the partitioning of NC cells into subpopulations with distinct migration routes and potencies^1^. The evolution of these developmental modules is poorly understood. Endothelin (Edn) signaling is unique to vertebrates, and performs various functions in different NC subpopulations^2–5^. To better understand the evolution of NC patterning, we used CRISPR/Cas9-driven mutagenesis to disrupt Edn receptors, ligands, and Dlx transcription factors in the sea lamprey, *Petromyzon marinus*. Lampreys and modern gnathostomes last shared a common ancestor 500 million years ago^6^. Thus, comparisons between the two groups can identify deeply conserved and divergent features of vertebrate development. Using *Xenopus laevis* to facilitate side-by-side analyses, we show here that lamprey and gnathostomes display fundamental differences in Edn signaling function. Unlike gnathostomes, both lamprey Ednrs cooperate during oropharyngeal skeleton development. Furthermore, neither paralog regulates *hand* transcription factors, which are required for mandible development in gnathostomes. We also identify conserved roles for Edn signaling in *dlx* gene regulation, pigment cell, and heart development. Together our results illustrate the stepwise neofunctionalization and specialization of this vertebrate-specific signaling pathway, and suggest key intermediate stages in the early evolution of the NC.

## Introduction

The proper migration, patterning, and differentiation of most NC subpopulations requires Edn signaling. In model gnathostomes, Edn ligands secreted by oropharyngeal epithelia are bound by Edn receptors (Ednrs) expressed by NC cells. In zebrafish and mouse, disruption of *edn1* or *edn* receptor A (*ednra*) results in a hypomorphic pharyngeal skeleton, skeletal element fusions, and ventral-to-dorsal transformations of oropharyngeal cartilages and bones^2,3,7–10^. In zebrafish, the increased dorsoventral symmetry and lack of a jaw joint causes a *‘sucker’* phenotype reminiscent of modern agnathans^2^. In both mouse and zebrafish, the skeletal phenotype of *edn1/ednra* mutants appears due to reduced expression of *dlx* and *hand* genes in cranial NC cells^3,11^. In non-skeletogenic NC, loss of *edn3* or *ednrb* causes aberrant migration and/or loss of pigment cells^5,12,13^. In mammals, these defects are accompanied by disruptions in NC-derived enteric neuron development^14^.

Lamprey expresses *edn, ednr, dlx*, and *hand* in patterns similar to their gnathostome cognates^15–17^ though lamprey and gnathostome NC derivatives differ substantially. In addition to lacking jaws, the lamprey oral skeleton consists of a specialized pumping organ made of a chondroid tissue called mucocartilage^18–20^ (Fig. 1A, Supplemental Fig. 1A and B). In the posterior pharynx, the branchial skeleton is a network of cell-rich hyaline cartilage bars, and a ventral mass of mucocartilage^18–20^. In the trunk, the lamprey PNS lacks sympathetic ganglia, Schwann cells, and vagal NC-derived enteric neurons^18^. These differences, and the unclear phylogenetic relationships between gnathostome and lamprey *edn* and *dlx* homologs, have led to speculation that these genes acquired new roles in patterning the NC of stem gnathostomes^17,21^.

**Figure 1.**
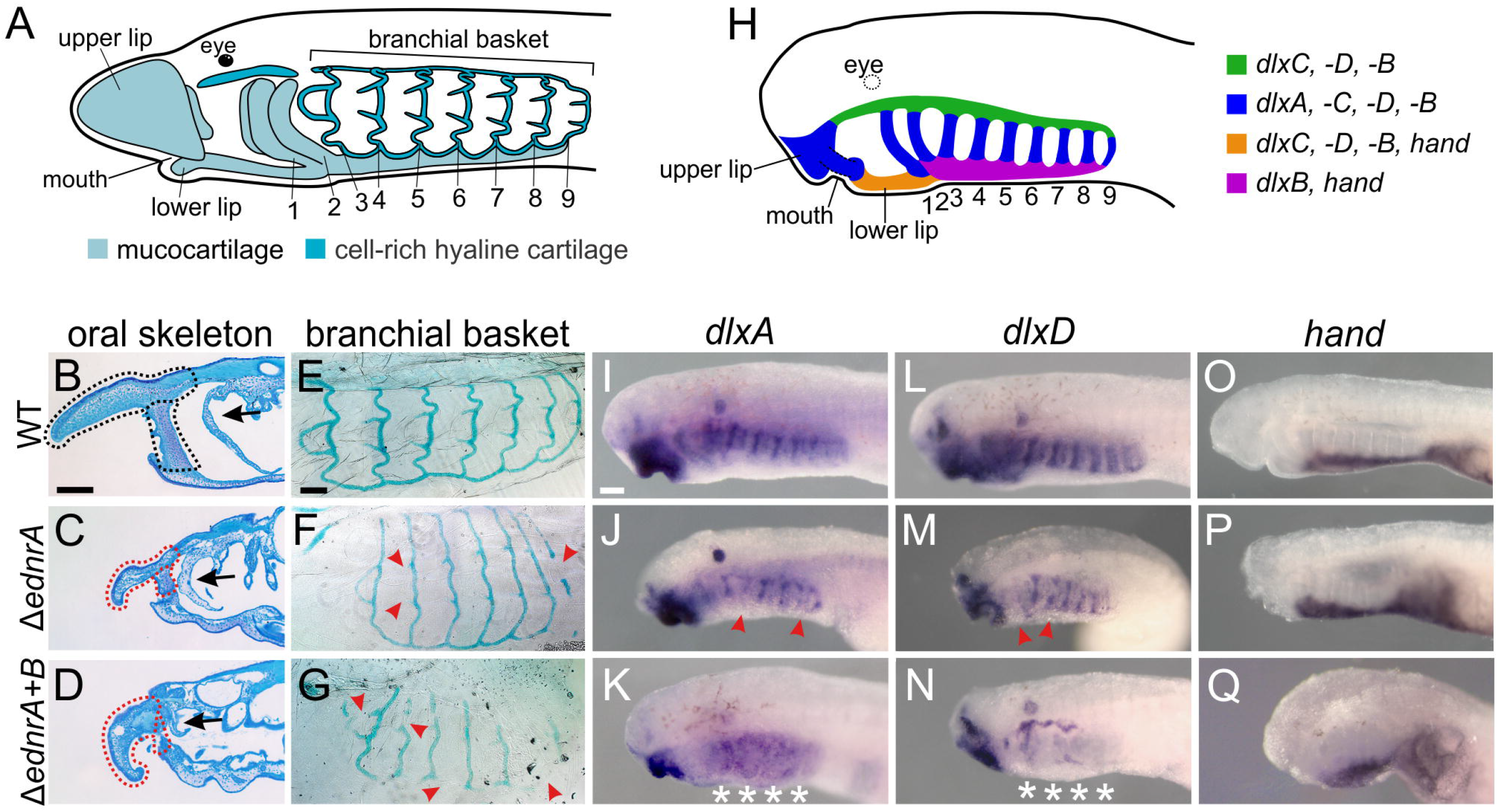
Lamprey Δ*ednr* larvae have pharyngeal skeleton defects, and reduced intermediate domain dlx expression. **A**, Illustration of the larval sea lamprey pharyngeal skeleton at st. T30. **B-G**, Toluidine blue-stained sagittal section of the oral mucocartilage (B-D) and flat-mounted alcian blue stain of the branchial basket (E-G) at. st. T30 in wildtype (B,E) Δ*ednra* (C,F) and Δ*ednra+b* (D,G) larvae. Δ*ednra* and Δ*ednra+b* mutants have reduced pharyngeal skeletons with gaps and missing elements (arrows). **H**, Illustration of *dlx* and *hand* expression in the lamprey head at T26.5 **I-Q**, *dlxA, dlxD*, and *hand* expression in WT (I,L,O), Δ*ednra* (J,M,P), and Δ*ednra+b* (K,N,Q) mutants. Disruptions in the differentiated pharyngeal skeleton at. st. T30, and intermediate domain *dlx* expression at st. T26.5 are seen in both Δ*ednra* and Δ*ednra+b* larvae, but are most pronounced in Δ*ednra+b* individuals. In contrast, ventral *hand* expression persists in Δ*ednra* and Δ*ednra+b* larvae. Asterisks indicate pharyngeal arches with missing expression. All scale bars indicate 100 μm. Scale bar in B applies to B,C,D. Scale bar in E applies to E,F,G. Scale bar in I applies to I-Q. All panels show left lateral views. See Supplemental Tab. 1-4 for sgRNA sequences, genotyping data, numbers affected, and statistical analyses.

## Results and Discussion

To test this idea, we optimized a method for efficient Cas9-mediated mutagenesis in the sea lamprey^22^ and used it to disrupt the function of *ednrs, edns*, and *dlxs*. Recent assembly of the sea lamprey germline genome^23^ supports previous reports that lamprey has one *ednra* and one *ednrb*, and six *edns, ednA-F*^17,24^. Because multiple, possibly redundant *edns* are coexpressed in lamprey embryos, we first mutagenized lamprey *ednrs*. Targeting two unique protein-coding sequences to control for off-target effects (Supplemental Tab. 1), we found that Cas9-mediated F0 mutation of *ednra* (Δ*ednra*) resulted in a hypomorphic pharyngeal skeleton with gaps in the branchial basket, ectopic melanophores, and heart edema (Fig. 1B,C,E,F, Supplemental Figs. 1C, 2, and 3). While the Δ*ednra* phenotype resembles gnathostome *ednra/edn1* mutants, including the ectopic pigment cells^25^, it differs from the reported effects of an Edn signaling inhibitor^26^, likely reflecting the specificity of CRISPR/Cas9.

In gnathostomes, Ednra/Edn1 signaling acts, in part, by regulating the expression of *dlx* paralogs in the intermediate pharynx and *hand* genes in the ventral pharynx^3,11,27^. We asked if lamprey *ednra* regulates these genes in lamprey NC cells. Despite divergent histories of *dlx* duplication and loss^28^, lamprey Δ*ednra* larvae exhibited reduced *dlx* expression in the intermediate domain (Fig. 1H,I,J,L,M, Supplemental Fig. 4) though ventral *hand* expression persisted (Fig. 1O and P). To ensure that this difference was not due to the mosaicism of Cas9-mediated F0 mutagenesis, we created *Δednra* and *Δedn1 Xenopus laevis* larvae. As in other gnathostomes, we observed a hypomorphic pharyngeal skeleton and a loss of the jaw joint that correlated to a loss of intermediate *dlx*, and ventral *hand*, expression (Supplemental Fig. 5). These data suggest Edn-dependent intermediate domain expression of *dlx* was likely present in the lamprey/gnathostome common ancestor, while Edn-dependent expression of *hand* is unique to gnathostomes.

Sea lamprey *ednrs* are broadly coexpressed in skeletogenic NC, a pattern not observed in any gnathostome, though some gnathostomes briefly express *ednrb1* in these cells^24^. This raised the possibility that *ednrb* and *ednra* both function in lamprey pharyngeal skeleton development. We thus used three separate sgRNAs to mutagenize *ednrb* alone (Δ*ednrb*), and together with *ednra* (Δ*ednra+b*). We found that, like gnathostome *ednrb/edn3* mutants, lamprey *Δednrb* and Δ*ednra+b* individuals have severe reductions in melanophores, the only pigment cells discernible in lab-raised larvae (Supplemental Fig. 6). However, unlike any reported gnathostome *ednrb* mutants, alcian blue staining revealed mild skeletal defects in Δ*ednrb* larvae (Supplemental Figs. 1D, 6B). Furthermore, Δ*ednra+b* larvae had losses and reductions of skeletal elements that were more severe than either single mutant (Fig. 1D and G, Supplemental Figs. 1E and 6C). Intermediate *dlx* expression was also more reduced in *Δednra+b* larvae than in Δ*ednra* or Δ*ednrb* larvae, though Δ*ednrb* individuals showed no apparent *dlx* reduction (Fig.1K and N, Supplemental Fig. 4). In contrast, expression of *hand* in the ventral domain remained in Δ*ednrb* and Δ*ednra+b* individuals (Fig. 1Q, Supplemental Fig. 6). These results demonstrate cooperative roles for lamprey *ednra* and *ednrb* in lamprey skeletogenic NC, and confirm divergence of ventral pharyngeal NC specification between gnathostomes and lamprey.

To better understand the function of Edn signaling in lamprey NC, we analyzed the expression of several cranial NC markers in Δ*ednra*, Δ*ednrb*, and Δ*ednra+b* embryos and larvae. *twistA*^29^ and *soxE2*^30^ expression at st. T22-23 in *Δednra+b* embryos suggests that the specification and initial migration of cranial NC cells is mostly normal in Δ*ednr* individuals (Fig. 2A and B). In T26.5 Δ*ednra* and Δ*ednrb* larvae, *myc*^29^, *ID*^31^, *soxE1*^32^, *twistA*^29^, and *msxB*^15^ expression NC also persisted in most postmigratory NC, further indicating largely normal cranial NC development, though subtle migration defects are also possible (Fig. 2C and D, Supplemental Fig. 7). In contrast, at T26.5, both Δ*ednra* and Δ*ednrb* larvae displayed clear reductions in *soxE2*^30^ expression in the forming branchial bars and *lecticanA*, an *aggrecan* homolog, in the branchial bars and differentiating mucocartilage (Fig. 2E-G,I-K, Supplemental Fig. 1G-K), though with different penetrance (Supplemental Tab. 2). Similar reductions in *soxE2* and *lecticanA* were observed in Δ*ednra+b* larvae, which also displayed reductions in *twistA* and *soxE1*, and localized loss of *ID* transcripts in oral mucocartilage precursors (Supplemental Fig. 7). Taken together, our results show that mutation of either, or both, *ednrs* results in reduced of *soxE* expression and skeletogenic NC differentiation, with the disruptions occurring most consistently in Δ*ednra* and Δ*ednra+b* individuals (Supplemental Tab. 2). Reductions in skeletogenic NC are also seen following *edn1/ednra* perturbation in model gnathostomes^8,33,34^, though disrupted *soxE* (*sox9a*) expression has only been reported in zebrafish^33^. We thus visualized *sox9.S* expression in *X. laevis* Δ*ednra* larvae, and found reduced expression (Fig. 2O and p). This suggests regulation of *soxE* expression and NC skeletogenesis are deeply conserved functions of Edn signaling in vertebrates.

**Figure 2.**
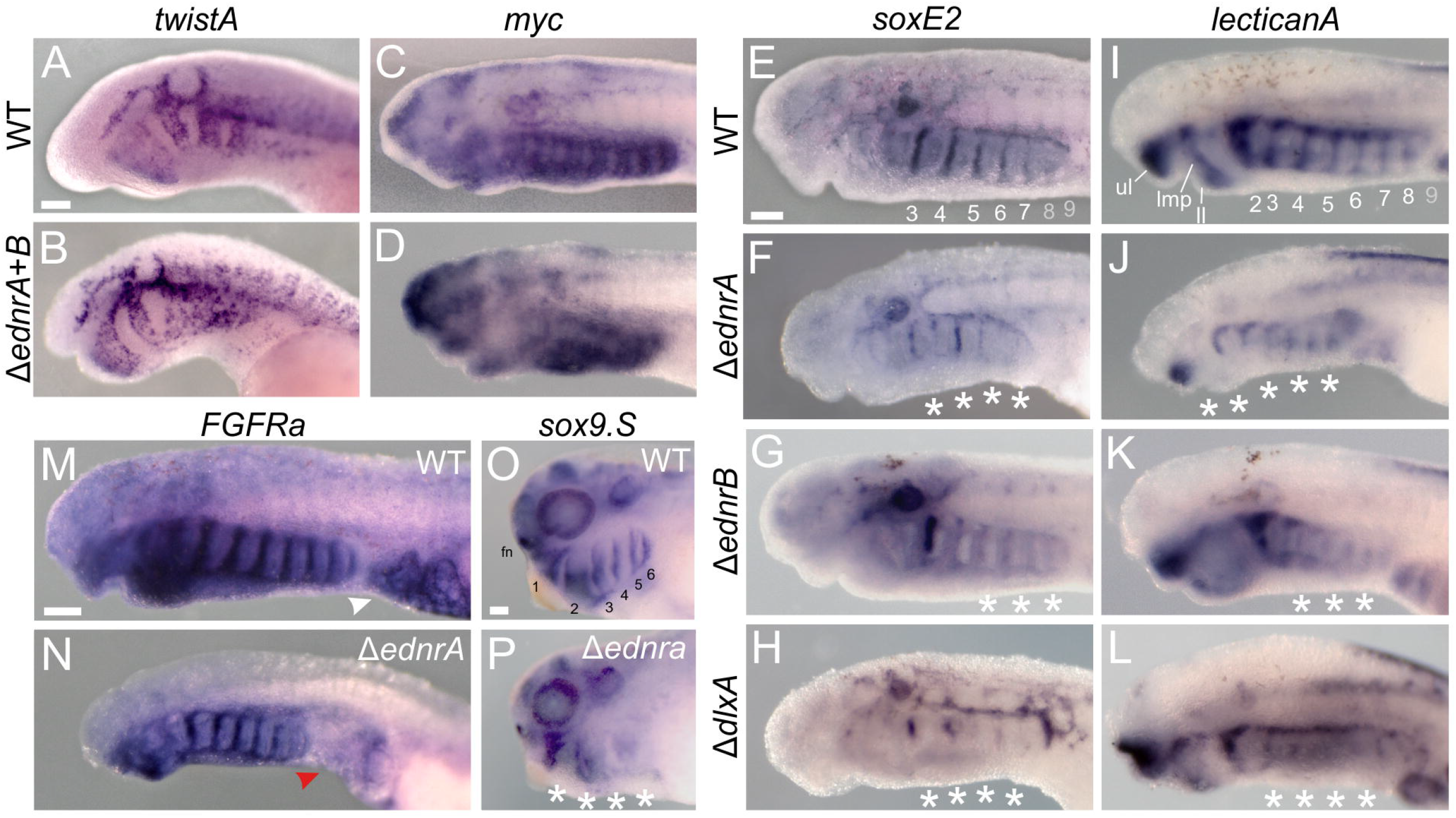
Skeletogenic NC development is disrupted in lamprey Δ*ednr* and Δdlx, and in *X. laevis Δednra* larvae, and cardiac mesoderm development is disrupted in lamprey Δ*ednra* larvae. **A-B**, Expression of *twistA* in migratory NC at st. T23 in WT (A) and Δ*ednra+b* larvae (B). **C-D**, Expression of *myc* in post-migratory NC at st. T26.5 in WT (C) and Δ*ednra+b* larvae (D). The expression of *twistA, myc*, and other general cranial NC markers (Supplemental Fig. 7), suggests that cranial NC formation is largely normal in Δ*ednr* larvae. **E-L**, reduced and discontiguous expression of *soxE2* (E-H) and *lecticanA* (I-L), in WT (E,I), Δ*ednra* (F,J), Δ*ednra+b* (G,K), and Δ*dlxA* (H,L) larvae at st. T26.5. Δ*ednra, Δednrb*, and Δ*dlx* mutants have similar disruptions in *soxE2* and *lecticanA* expression, showing skeletogenic NC development is disrupted when these genes are mutated. **M-N**, *FGFRa* expression in cardiac mesoderm (M) is reduced in Δ*ednra* mutants (N). **O-P**, Like lamprey Δ*ednra and Δednrb* larvae, *Xenopus laevis Δednra* larvae (N) have disruptions in *soxE* (sox9.s) expression in post-migratory NC (st. N.F. 33/34). All panels show left lateral views. All scale bars indicate 100 μm. Scale bar in A applies to A-D. Scale bar in E applies to E-L. Scale bar in M applies to M-N. Scale bar in O applies to OP. See Supplemental Tab. 1-4 for sgRNA sequences, genotyping data, numbers affected, and statistical analyses.

We next asked if *dlx* genes are effectors of Edn signaling in the lamprey pharyngeal skeleton by comparing the phenotype of Δ*ednr* and Δ*dlx* individuals. Mutation of *dlxA, dlxC*, and *dlxD* alone, or in combination, resulted in disruptions of *soxE2* and *lecticanA* expression similar to Δ*ednr* larvae (Fig. 2H and L, Supplemental Fig. 8A). At T30, Δ*dlx* individuals also had hypomorphic pharyngeal skeletons with gaps in the branchial basket, though they lacked the heart and pigment defects seen in Δ*ednr* larvae (Supplemental Fig. 8B). The similarity of Δ*ednr* and Δ*dlx* individuals suggest that Edn signaling works though *dlx* genes in lamprey skeletogenic NC.

Like mouse *ednra*^35^, lamprey *ednra* transcripts mark presumptive cardiac mesoderm^24^ and lamprey Δ*ednra* larvae have severe heart defects (Supplemental Figs. 2, 3). We thus examined the expression of the *FGFR* homolog, *fgfra*, in Δ*ednra* larvae. In addition to being transcribed in lamprey cardiac mesoderm, functional studies suggest *fgfra* signaling is required for lamprey heart development^22,36^. We observed a strong reduction in cardiac *fgfra* expression in Δ*ednra* individuals (Fig. 2M and N). This indicates that the heart edema seen in lamprey Δ*ednra* larvae is likely caused by reduced FGFR signaling in cardiac mesoderm.

Unlike *ednra, ednrb* is strongly expressed in trunk NC cells destined to form the dorsal root ganglia^24^. We thus examined the expression of several PNS markers^37–40^ in Δ*ednrb* and Δ*ednra+b* embryos and larvae (Fig. 3A-M). Despite the reduced heads of Δ*ednra+b* larvae, all major NC-derived PNS elements were present, including recently described SCPs^38^, though chromaffin-like cells near the pronephros were missing^41^ (Fig. 3A-C, arrowheads). This relatively mild defect is similar to gnathostome *edn3/ednrb* mutants and morphants which have only minor PNS deficiencies. These results suggest that development of the NC-derived PNS was largely Edn-independent in the last common ancestor of lamprey and gnathostomes.

**Figure 3.**
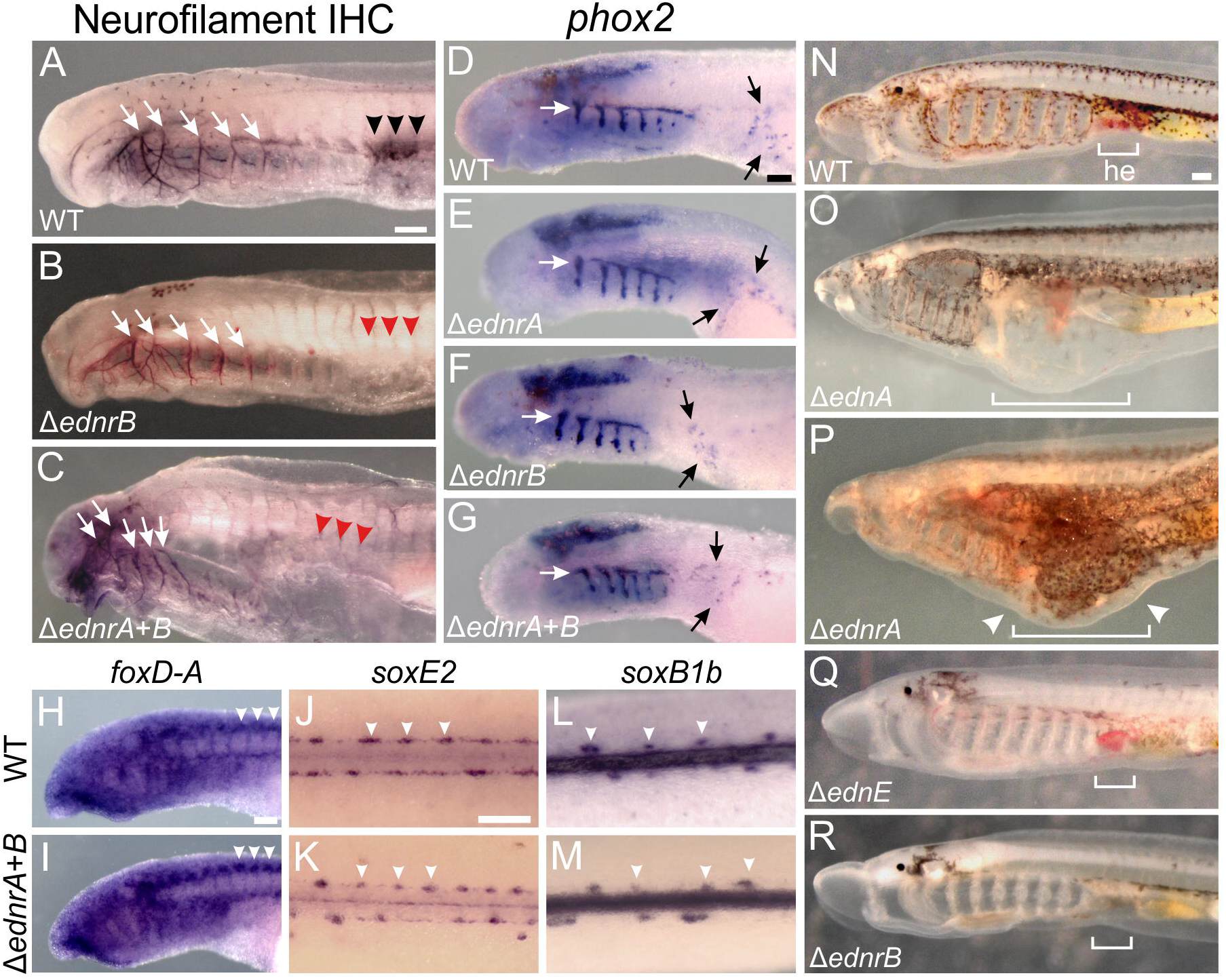
Most PNS elements form normally in lamprey Δ*ednr* larvae, and Δ*edn* and Δednr larvae have similar phenotypes. **A-C**, Neurofilament immunohistochemistry at st. T27 reveals all major facial nerves (white arrows) in WT (A), Δ*ednrb* (B), and Δ*ednra+b* mutants (C), though a population of Neurofilament-positive cells near the pronephros is missing (arrowheads B,C). **D-G**, *phox2* expression at st. 26.5 reveals forming epibranchial ganglia (white arrows) and the recently described “Schwann cell precursors” (black arrows) in WT (D), Δ*ednra* (E), Δ*ednrb* (F), *and Δednra+b* mutants G). Despite reduced heads, Δednr mutants have largely normal epibranchial ganglia. **H-M**, Comparisons of *foxD-A* (H,I), *soxE2* (J,K), and *soxB1b* (L,M) expression in WT (H,J,L) and Δ*ednra+b* mutants (I,K,M) at st. 26.5 suggest the dorsal root ganglia (white arrowheads) form normally in Δ*ednra, Δednrb*, and Δ*ednra+b* mutants. **N**, WT st. T30 larvae, **O-R**, Δ*edn* mutants phenocopy mild Δ*ednr* mutants. Δ*ednA* mutants (O) recapitulate the hypomorphic head and heart edema (brackets) of Δ*ednra* mutants (P), but lack the ectopic pigmentation caused by *ednra* disruption (arrowheads). Δ*ednE* mutants (Q) have reduced pigmentation, resembling Δ*ednrb* mutants (R). All scale bars indicate 100 μm. Scale bar in A applies to A-C. Scale bar in D applies to D-G. Scale bar in H applies to H-I. Scale bar in J applies to J-M. Scale bar in N applies to O-S. All panels show left lateral views, save J-M which are dorsal views of the trunk (anterior to left). See Supplemental Tab. 1-4 for sgRNA sequences, genotyping data, numbers affected, and statistical analyses.

*In vitro* binding assays^42^, and the similarity of *edn* and *ednr* mutant phenotypes^2,33^, suggest that Edn1 is the main ligand for Ednra, while Edn3 is the main ligand for Ednrb. To test if lamprey Ednra and Ednrb also have dedicated ligands, we mutated *ednA, ednC*, and *ednE*, the only *edns* expressed in tissue-specific patterns during lamprey development. Targeting *ednC* with three different sgRNAs yielded no reproducible mutant phenotype (see Methods). In contrast, lamprey Δ*ednA* larvae displayed a combination of heart edema and skeletal defects that resembled hypomorphic Δ*ednra* individuals (though without ectopic pigmentation), while Δ*ednE* larvae resembled Δ*ednrb* larvae (Fig 3N-R, Supplemental Fig. 9). The incomplete loss of pigment in Δ*ednE*/Δ*ednrb* larvae mimics gnathostome *edn3/ednrb* mutants^43^, though the variable phenotype could also reflect the mosaicism inherent in F0 mutagenesis. To address this, we created Δ*edn3 X. laevis*. We found a high percentage had a complete loss of NC-derived pigmentation (Supplemental Fig. 10, Supplemental Tab. 1), like *edn3* mutant salamanders^44^, confirming that F0 mutagenesis can generate near-null *edn* phenotypes a high frequencies. We conclude that all modern vertebrates have an *edn* dedicated to *ednrb*, and that amphibian NC-derived pigment cell development appears particularly dependent on *edn3/ednrb* signaling.

Inconclusive phylogenies^17,24,45^, similar expression patterns^24^, and similar mutant phenoptypes suggest that lamprey *ednA* and *ednE* could be cryptic orthologs of gnathostome *edn1* and *edn3*, respectively. We thus used the recently completed sea lamprey germline genome^23^ to reevaluate *ednr* and *edn* phylogeny. Conserved synteny strongly confirms orthology of lamprey and gnathostome *ednras* and *ednrbs^24,46^* (Supplemental Figs. 11,12). In contrast, phylogenetic analysis of flanking genes indicate that *ednA* and *ednE* are likely part of the *edn2/4 and edn1/3* paralogy groups, respectively (Supplemental Figs. 11,12). Thus, while an Edn largely dedicated to Ednrb likely arose before the divergence of lamprey and gnathostomes, the lack of orthology between *edn1* and *ednA* reveals a mainly Ednra-specific ligand evolved at least twice.

Work in invertebrate chordates suggests that the NC likely evolved from a population of migratory CNS cells that generated neurons and/or pigment cells^47,48^. In contrast, the NC of modern vertebrates is a patterned tissue made of many distinct subpopulations. Our work supports a model for the stepwise evolution of the major NC subpopulations (Fig. 4). We show here that lamprey larvae lacking all, or most, Edn signaling have a largely normal PNS, but an oropharyngeal skeleton reduced to unconnected cartilage bars. The similarity of this phenotype to the presumed vertebrate ancestor suggests Edn signaling evolved in an early stem vertebrate that had all major NC derivatives^49–52^. Because both *ednrs* are needed for normal oropharyngeal skeleton development in lamprey, and some gnathostomes express *ednrb* in skeletogenic NC^24^, we further posit the ancestral pre-duplication Ednr functioned in this derivative. After *ednr* duplication, but before the evolution of jaws, the subfunctionalization and specialization of *ednra* and *ednrb* contributed to the developmental divergence of the pigment and skeletogenic NC lineages. Finally, differences in the regulation of *hand*, and the role of *edn3/ednrb* in ENS development, suggest changes in Edn signaling targets may underlie divergence of the lamprey and gnathostome oropharyngeal skeletons and PNS.

**Figure 4.**
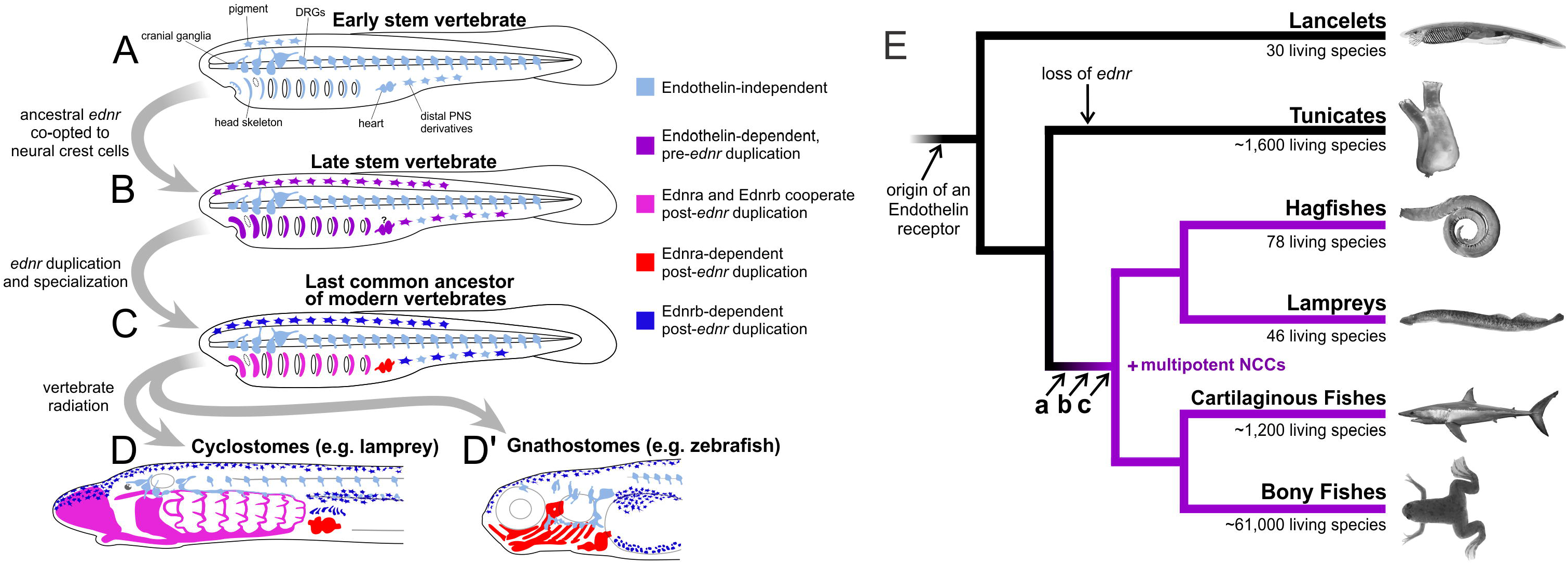
The stepwise evolution of Edn signaling in neural crest development deduced from lamprey and gnathostome Edn/Ednr mutant phenotypes. **A**, The similarity of Edn signaling-deficient lamprey larvae to basal vertebrate fossils, and the finding that Edn signaling is dispensable for early NC development in all modern vertebrates, suggest the vertebrate ancestor had a fully potent NC that developed in the absence of Edn signaling. **B**, The large reductions in NC-derived skeletal tissue and pigment cells in lamprey and gnathostome Δ*ednra+b* mutants, but not NC-derived peripheral neurons, suggest the recruitment of Endothelin signaling contributed to the expansion of skeletal, and other non-neural NC derivatives, in stem vertebrates. **C**, Subsequent duplication and specialization of the *ednra* and *ednrb* signaling pathways resulted in three NC subpopulations with different Endothelin signaling requirements. Differences in the phenotypes of lamprey and gnathostome Edn signaling mutants suggest the roles of Edn signaling in pharyngeal skeleton patterning has diverged in jawed and jawless vertebrates. **D**, In the sea lamprey, Ednrb works with Ednra in skeletogenic NC, while *hand* transcription in the ventral pharynx is Endothelin signaling-independent. **D’**, In modern gnathostomes, pharyngeal skeleton development is Ednrb-signaling independent, and *hand* expression in the ventral pharynx requires Ednra signals. **E**, The deduced character states in A-C mapped onto a phylogenetic tree of extant chordate groups.

## Methods

### “F0” mutagenesis strategy

We used CRISPR/Cas9-mediated mutagenesis to induce deletions and insertions (indels) into the protein-coding exons of injected “F0” sea lamprey (*Petromyzon marinus*) and African clawed frog (*Xenopus laevis*) embryos as previously described^22,53–57^. Though CRISPR/Cas9 is highly efficient in sea lamprey, differences in the efficiency of individual sgRNAs results in different ratios of wildtype and mutant alleles in F0 mosaic mutants. This variable mosaicism results in different sgRNAs producing phenotypically mutant individuals at different frequencies, with a range of severities. Previous work shows targeting an evolutionarily conserved, embryonically-expressed gene typically results in 20-90% of injected individuals displaying a gene-specific mutant phenotype^22,53–56^. Work in our lab with 35 guides targeting 20 different developmental regulators confirms this, with an average of 46% phenotypically mutant individuals produced per gene-specific sgRNA (Supplemental Tab. 1, Supplemental Fig. 13).

Also as previously reported, the severity of a CRISPR/Cas9-induced phenotype correlates well with the percentage mutant alleles; with most “severely affected” F0 mosaic mutants typically exhibiting 75%-100% mutant (indel) alleles^22,53–56^. Consistent with this, the 74 severely affected phenotypic mutants selected for genotyping in this study had an average of 88% indel alleles at targeted loci (Supplemental Tab. 4). Importantly, every severely affected individual selected for genotyping had indel mutations at the targeted locus. Thus, as with traditional inbred mutant lines, the phenotype of CRISPR/Cas9-generated F0 mosaic mutants is a strong predictor of their genotype.

Based on these observations, we devised a strategy for creating, selecting, and analyzing CRISPR/Cas9-generated sea lamprey and *X. laevis* mutants. First, two or more unique sgRNAs were designed against protein-coding exons of the gene of interest. When possible, we selected unique, but evolutionarily conserved regions to increase the chances that in-frame deletions will disrupt functionally critical domains and yield loss-of-function alleles. Second, individual sgRNAs were co-injected with Cas9 protein or mRNA into zygotes or, in the case of *X. laevis*, zygotes and two-cell stage embryos. Third, F0 injected embryos were monitored daily and scored for morphological defects. Fourth, morphological defects associated with two or more sgRNAs targeting the same gene were designated as the “mutant phenotype” for that gene. For example, the unique pigmentation defect seen when targeting *ednrb* exons was deemed the “*ednrb* mutant phenotype” only after two different sgRNAs targeting the *ednrb* locus produced the same defect. Fifth, mutagenesis of the targeted loci was confirmed by genotyping several representative severely affected phenotypic mutants (see below for genotyping method). Sixth, once mutant genotype and mutant phenotype were linked by showing all selected mutants had mutant alleles, severely affected phenotypic mutants were picked for analyses via *in situ* hybridization, alcian blue staining, immunohistochemistry, and toluidine blue staining (see below for protocols). For *dlx* sgRNAs, which resulted in unusually high mortality before larval stages, likely due to the early function of *dlx* genes in neurectoderm patterning, severe phenotypic mutants were lightly fixed and genotyped after *in situ* hybridization analysis as recently described^54,55^. This additional step was performed to reconfirm the link between mutant phenotype and mutant genotype in the relatively small number of surviving *dlx* mosaic mutants.

### *P. marinus* sgRNA/Cas9 injections

We mutagenized the *P. marinus dlxA, dlxC, dlxD, ednA, ednC, ednE, ednra*, and *ednrb* loci by injecting zygotes with at least two unique sgRNAs per gene (Supplemental Tab. 1). To create *ednra+b* double mutants, zygotes were injected with four different combinations of the two most effective *ednra* and *ednrb* guides. *dlxA+C+D* triple mutants were created using a single sgRNA 100% complementary to *dlxA* and *dlxD*, with one mismatch to *dlxC* (Supplemental Tab.1). As previously described, sgRNA target sites were chosen using all available transcriptome sequence data to avoid protein-coding off-targets^22^. Briefly, candidate sgRNA sequences demonstrating off-target matches with >80% overall identity in the target site, and >90% identity in the 3’ half of the target site (closest to the PAM site) to any off-target sequence (with an NGG PAM site) were not used. Lamprey zygotes were injected as previously described with approximately 5 nL of a solution containing 400 pg of sgRNA, and either 800 pg of Cas9 protein (a 2:1 ratio of protein:sgRNA by mass) or 1 ng of Cas9 mRNA, 5 mg/mL lysinated rhodamine dextran (LRD), and nuclease free water. For *ednra* + *ednrb* combined experiments, 200 pg of each of two sgRNAs were used with 800 pg of Cas9 protein. Approximately 200-500 zygotes were injected per experiment, and each sgRNA/Cas9 combination was injected into zygotes from at least two different pairs of wild caught sea lampreys.

As in other vertebrates^58^, microinjection of lamprey embryos causes increased mortality before gastrulation and developmental delay compared to uninjected sibling controls^22^. Due to differences in female health, person injecting, and progression of the spawning season, this microinjection-induced mortality can range from 10-90%. However, after gastrulation, clutches of microinjected embryos have a survivorship to early larval stages (T26-T30) similar to uninjected siblings, typically around 90%. This was true for all sgRNAs tested in this study, except for the *dlx* sgRNAs, which had substantially increased mortality to larval stages compared to uninjected siblings, resulting in 30-40% survival to T26.5. We suspect this is due to the early roles of *dlx* genes in neurectoderm patterning.

### *X. laevis* sgRNA/Cas9 injections

Both the “Long” (-*L*) and “Short” (-*S*) homoeologs of *X. laevis edn1, edn3*, and *ednra* were simultaneously targeted^57^ (Supplemental Tab. 1). Zygotes or two-cell embryos were injected with a 5-10 nL droplet containing 800 pg of a single sgRNA targeting both *edn3.L* (formerly *edn3-a*) and *edn3.S* (-b), or 400 pg of sgRNAs targeting *edn1* and *ednra*, and either 1 ng of Cas9 mRNA, or 1.6 ng of Cas9 protein. Approximately 50-200 zygotes were injected per experiment.

### *P. marinus* CRISPR/Cas9 controls

To demonstrate that the phenotypes associated with each sgRNA injected were due to disruption of the targeted genes, rather than to off-targets, each *P. marinus* gene was targeted with at least two unique sgRNAs. All sgRNAs targeting the same gene produced the same mutant phenotype, though usually with different efficiencies (Supplemental Tab. 1).

To further validate sgRNA specificity in *P. marinus*, and to ensure that the CRISPR/Cas9 method does not artefactually cause any of the described defects, we used two negative control strategies. In addition to the negative control sgRNA described in our methods paper^22^ we tested an intron-spanning sgRNA partially complementary to two separate exons of the *P. marinus ednrb* gene (see Supplemental Tab. 1 for sequence). Neither sgRNA produced a phenotype (Supplemental Fig. 13), though both resulted in a slight developmental delay, as previously reported^22^. In addition to these “untargeted” sgRNA negative controls, we also injected more than 20 other sgRNAs complementary to the exons other *P. marinus* developmental genes (Supplemental Fig. 13). These sgRNAs were designed to disrupt developmental regulators expressed in the developing head at the same time as *ednr, edn*, and *dlx*. None of these negative control sgRNAs yielded the *ednr or edn* mutant phenotypes, though three sgRNAs (a2cg1, p19g1, and w11g3) produced phenotypes grossly similar to *dlx* mutants (Supplemental Fig. 13).

Severe heart edema (approximate heart volume greater than 3x normal by visual inspection) is part of both the *ednra* and *ednA* mutant phenotypes, and occurs at a high frequency in embryos injected with sgRNAs targeting *fgf8/17/18*^22^ (Supplemental Fig. 13). This raised the possibility that heart edema could be a non-specific side-effect of sgRNA/Cas9 injection. To test this, we counted the number of negative control larvae, aside from those injected with *fgf8/17/18* sgRNA, displaying heart edema (Supplemental Fig. 13). Of 21 pools of larvae injected with 21 different negative control sgRNAs, 9 pools displayed no detectable heart edema, while 11 displayed heart edema of various severities at a frequency of 7.7% or lower. 1 sgRNA yielded severe heart edema at a frequency of 27%. These data show that severe heart edema is not a general side-effect of the CRISPR/Cas9 method in lamprey.

### *X. laevis* CRISPR/Cas9 controls

An *edn3* morphant phenotype was previously reported in *X. laevis*^59^. An sgRNA designed to simultaneously target the *edn3.L* and *edn3.S* homoeologs yielded a severe version of the *X. laevis edn3* morphant phenotype that mimicked salamander *edn3* mutants^44^, confirming its specificity. For *edn1* and *ednra*, we designed separate sgRNAs against the *L* and S homoeologs and performed negative controls by individually injecting each sgRNA separately as reported previously^57^. This strategy relies on redundancy of the *X. laevis* homoeologs to show that neither sgRNA alone causes any spurious morphological defects. The fact that defects are only obtained by simultaneous disruption of homoeologs, serves as a control showing that the phenotype is specifically due to a loss of *edn1* and *ednra* function^57^.

### *P. marinus* husbandry

*P. marinus* fertilizations and husbandry were carried out as described previously^22^. Adult spawning phase sea lampreys were housed in 200L tanks containing reverse osmosis purified water with 800-1000 ppm artificial sea salt. Water in the tanks was completely replaced daily. Once ripe, the animals were stripped of gametes into Pyrex dishes, where *in vitro* fertilization took place in deionized water containing 400-600 ppm artificial sea salt. All animals were wild-caught from fresh water streams during their late spring/early summer spawning season, with the majority being derived from an invasive population in Lake Huron. A small fraction (1%), were trapped at the Holyoke Dam in Massachusetts. Each sgRNA was injected into clutches from at least of two different pairs of adults. Embryos and larvae were kept at 18°C in Pyrex dishes containing deionized water and 400-600 ppm artificial sea salt. Depending on female health and time of year, uninjected sea lamprey embryos display survivorship to st. T26.5 from 1%-99%. Dead embryos and larvae were removed daily from each dish, and the water was changed at least every other day. All *P. marinus* staging was according to Tahara^60^. All *P. marinus* husbandry and experiments were in accordance with CU-Boulder IACUC protocol #2392.

### *X. laevis* husbandry

*X. laevis* fertilizations and husbandry were performed according to standard methods^61^. Adult females were induced to ovulate via injection of Human Chorionic Gonadotropin (HCG), and eggs were stripped into petri dishes. Testes were dissected from males, homogenized, and applied to the eggs for *in vito* fertilization. All frog staging was according to Nieuwkoop & Faber^61^. All *X. laevis* husbandry and experiments were in accordance with CU-Boulder IACUC protocol #2392.

### Scoring of mutant phenotypes

Successfully injected embryos were identified by fluorescence of the LRD lineage tracer at 4-6 days post fertilization and dead and LRD-negative larvae were discarded. Successfully injected embryos and larvae were then monitored for morphological abnormalities as they developed. Suites of morphological defects associated with injection of a particular sgRNA, and also seen when injecting one or more other sgRNAs targeting the same gene, were designated as the “mutant phenotype” for that gene. Of embryos and larvae displaying the “mutant phenotype”, we deduced, based on previous work, that most severe had more than 75% mutant alleles and were likely near null-mutants^22,53–56^. This was assumption was supported by genotyping representative severe mutants for all targeted genes (see “Genotyping” section below)

For each gene, we focused on “severely affected” mutants for detailed morphological and histological analyses. The severe mutant phenotype of all genes was apparent at pharyngula stages onward (st. T26.5 for lamprey, st. 41 for *X. laevis*) and defined as follows. For *X. laevis* Δ*edn1* and Δ*ednra* the severe mutant phenotype was defined as a reduction in head size (all structures anterior to the heart) to approximately 70% of WT size or smaller. For *P. marinus* Δ*ednA* the severe mutant phenotype was defined as a reduction in head size to approximately 70% of its WT size or smaller, together with heart edema. For Δ*ednra*, the severe mutant phenotype was defined as a reduction in head size to approximately 70% of its WT size or smaller, together with heart edema, and ectopic pigmentation around the heart. For Δ*dlxA*, Δ*dlxC*, and Δ*dlxD*, severe mutants were defined as having a head reduced to approximately 70% of WT size or smaller. For Δ*edn3*, Δ*ednE*, and Δ*ednrb* severe mutants were defined as having a 50% reduction in the number of melanophores or greater (in the case of *X. laevis* injected unilaterally at the 2 cell stage, this applies only to the injected side). For the Δ*ednra+b*, the severe mutant phenotype was defined as an approximately 70% reduction in head size, heart edema, and approximately 50% reduced pigmentation. All larvae demonstrating a “severe mutant phenotype” were counted and are presented as fraction of the total number of LRD- positive embryos and larvae that survived to fixation at a stage were phenotype could be scored (Supplemental Tab. 1, 2).

As in other vertebrates^58^, sea lamprey embryos injected with negative control sgRNAs, DNA constructs, or any other synthetic oligonucleotide, display a slight developmental delay. In sea lamprey we find that a delay of ~5% is typical, i.e. 10 day old injected embryos and larvae typically appear 9.5 days old compared to unmanipulated siblings. Thus, developmental events such as somite segregation, yolk absorption, gill openings, and melanin deposition^60^ were used, rather than days post-fertilization, to stage-match mutant and negative control embryos.

### Genotyping

To confirm successful mutagenesis, individual severe mutants were genotyped by preparing genomic DNA, PCR amplifying the target site, subcloning the amplicons, and Sanger sequencing individual alleles as previously described^22,53–56^. Target sites and genotyping primers for each sgRNA are in Supplemental Tab. 1. We genotyped at least 3 severely affected individuals for each targeted gene or combination of genes (Supplemental Fig. 2, 5, 6, 8-10, Supplemental Tab. 2) except in the case of *P. marinus ednrb* sgRNA2, which likely lies immediately adjacent to an intron/exon boundary conserved across jawed vertebrates (on the 5’ end of exon 4 in zebrafish ednraa [NM_001099445.2]), and is incompletely assembled in all three publicly available genomic assemblies (including the 2017 petMar3^23^). For *dlx* mutants, genotyping after histological analysis of lightly fixed mutants was performed as previously described^54–56^.

Frequently, we found six or more unique indel alleles at a given locus in a single specimen (in *X. laevis*, we consider the homoeologous”*L*” and “*S*” loci separately), which indicates that biallelic Cas9-driven mutagenesis is still occurring after the second cleavage event in both species. As previously reported^22,57,62^, when insertions of DNA fragments were discovered, these motifs often appeared on reverse or forward strand very close to the target site/lesion (see green and purple nucleotides in Supplemental Fig. 2, 5, 6, 8-10).

### Histological staining, *In situ* hybridization, and immunohistochemistry

All *in situ* hybridization (ISH), alcian blue cartilage staining, and toluidine blue staining was carried out as described previously^15,63,64^. Neurofilament immunohistochemistry (IHC) was as described previously^39^, with the addition of 1% dimethyl sulfoxide (DMSO) to the phosphate buffer solution prior to the blocking step. To ensure equivalent signal development in injected and WT individuals, morphologically stage-matched WT embryos and larvae were included in every ISH, IHC, and histological staining experiment, with WT and treated larvae kept in same tubes, with the caudal 1/4 cut-off for identification when necessary. In addition, to verify that none of the disrupted expression patterns or aberrant histology of mutants could be explained by slight developmental delay, a known side-effect of microinjection, WT embryos one stage younger were also used for comparisons, i.e. morphological st. T26.5 mutant larvae were compared to both morphological st. T25.5 and st. T26.5 wildtype larvae. The number of embryos and larvae processed for each histological method, and the frequencies of aberrations, are reported in Supplemental Tab. 2.

### Statistical analyses

We have never observed the *ednra, ednrb, ednA*, and *ednE* mutant phenotypes in wildtype or negative control embryos. In other words, the *ednra, ednrb, ednA*, and *ednE* phenotypes are only seen in embryos and larvae injected with Cas9 and sgRNAs targeting these genes. Similarly, we have never observed the reduced expression patterns we report in wildtype or negative control embryos. However, non-specific body axis deformities (mainly incomplete yolk-sac extension) occur at a frequency of 5-8% in surviving uninjected, and negative control-injected larvae. While these deformities are qualitatively different from the *ednra, ednrb, ednA*, and *ednE* mutant phenotypes, we used this background level of developmental deformity as a proxy to estimate mutant phenotype frequencies in negative control sea lamprey larvae (Supplemental Tab. 3). Using the conservative estimate that one out of ten negative control (untreated) individuals will spontaneously display the observed phenotypes, we applied Fisher’s exact test to evaluate the null hypothesis that our treatments can be explained by a high ‘background’ level of developmental deformities. This hypothesis is rejected with P values of <.02 for all mutant phenotypes, with the majority having P values <<.0001 (Supplemental Tab. 3).

Most *in situ* hybridization (ISH) assays were performed on embryos and larvae displaying the severe morphological phenotype. Because these specimens were non-randomly selected phenotypic mutants, statistical analysis is inappropriate. For pre-selected phenotypic mutants, we report the fraction of those assayed by ISH displaying disrupted gene expression patterns in Supplemental Tab. 3. The remaining ISH assays were performed on embryos before the mutant phenotype became apparent and severe mutants could be selected. In these cases, selected individuals were a random sample of the pool of sgRNA/Cas9 individuals and could be compared to untreated controls with Fisher’s exact test (Supplemental Tab. 3). For these experiments, we assumed spontaneous disruption of gene expression in 5 out of 100 of untreated, wildtype embryos and larvae. We view this assumption as conservative as we have never observed variation in gene expression patterns in wildtype embryos that have been properly processed for *in situ* hybridization or immunohistochemistry. Under this assumption, every reported effect of “no expression change” in this work is consistent with a null hypothesis of no effect/5% background levels of gene disruption (Fisher’s exact test P>0.35). For all genes we report as having discontiguous, missing, or otherwise reduced gene expression after treatment, the null hypothesis is rejected with P<<.0001 (Supplemental Tab. 3).

### Synteny and phylogenetic analysis

For the Endothelin receptors and ligands, we looked at synteny at each locus where possible (Supplemental Fig.11,12). The synteny analysis was performed by finding the coding sequences in the 2018 *P. marinus* genome^23^ via the UCSC genome browser (http://genome.ucsc.edu/) and comparing the neighboring genes to that of chicken (*Gallus gallus*) and/or human (*Homo sapiens*) as published previously^46^. For the Endothelin ligands, synteny information alone was ambiguous and amino acid similarity across large phylogenetic distances is poor, so we relied on the relatedness of the closely linked *hivep* and *phactr* gene products to deduce the likely evolutionary history of these gene families^45^ (Supplemental Fig. 12). For *Ednrs*, we repeated an amino acid similarity analysis according to the same methods as we used previously^24^, but with a subset of sequences. All phylogenetic and molecular evolutionary analyses were conducted using MEGA version 6^65^. See Supplemental Tab. 5. for all accession numbers associated with these analyses.

## Data Availability

All data generated or analyzed, and all methods used during this study are summarized in this published article (and its Supplemental information files). The raw data and images are available from the first and second authors upon reasonable request.

## Acknowledgements

The authors thank Scott Miehls at the USGS Hammond Bay Biological Station and Brittney Laflamme at the Holyoke Dam for providing adult sea lampreys, Bilge Birsoy, Jianli Shi, and Michael Klymkowsky for assistance with *X. laevis* fertilizations, Zachary Root for assistance with *X. laevis* and sea lamprey injections and husbandry, Claire Altier for assistance with juvenile *X. laevis* husbandry, and Shane Schwikert for *pro bono* statistics consultation. DMM, TAS, DJ, MR, JLM, and MVC were supported by National Science Foundation grants IOS 1656843, IOS 1257040, and IOS 0920751 to DMM. HPS and AWH were supported by the University of Colorado, Boulder Undergraduate Research Opportunities Program. DJ was supported by the European Union’s Horizon 2020 research and innovation programme under the Marie Skłodowska-Curie grant agreement No 751066 and by the Scientific Grant Agency of the Slovak Republic VEGA grant No.1/0415/17.

## Author Contributions

D.M.M. conceived the project. T.S., D.J., M.R., and D.M.M. designed the experiments. All authors performed experiments and collected data. D.M.M. and T.S. wrote the manuscript. All authors discussed and provided input on the final manuscript.

