## supplementary figures for "Cooption and Specialization of Endothelin Signaling Pathways Drove Elaboration of the Neural Crest in Early Vertebrates"

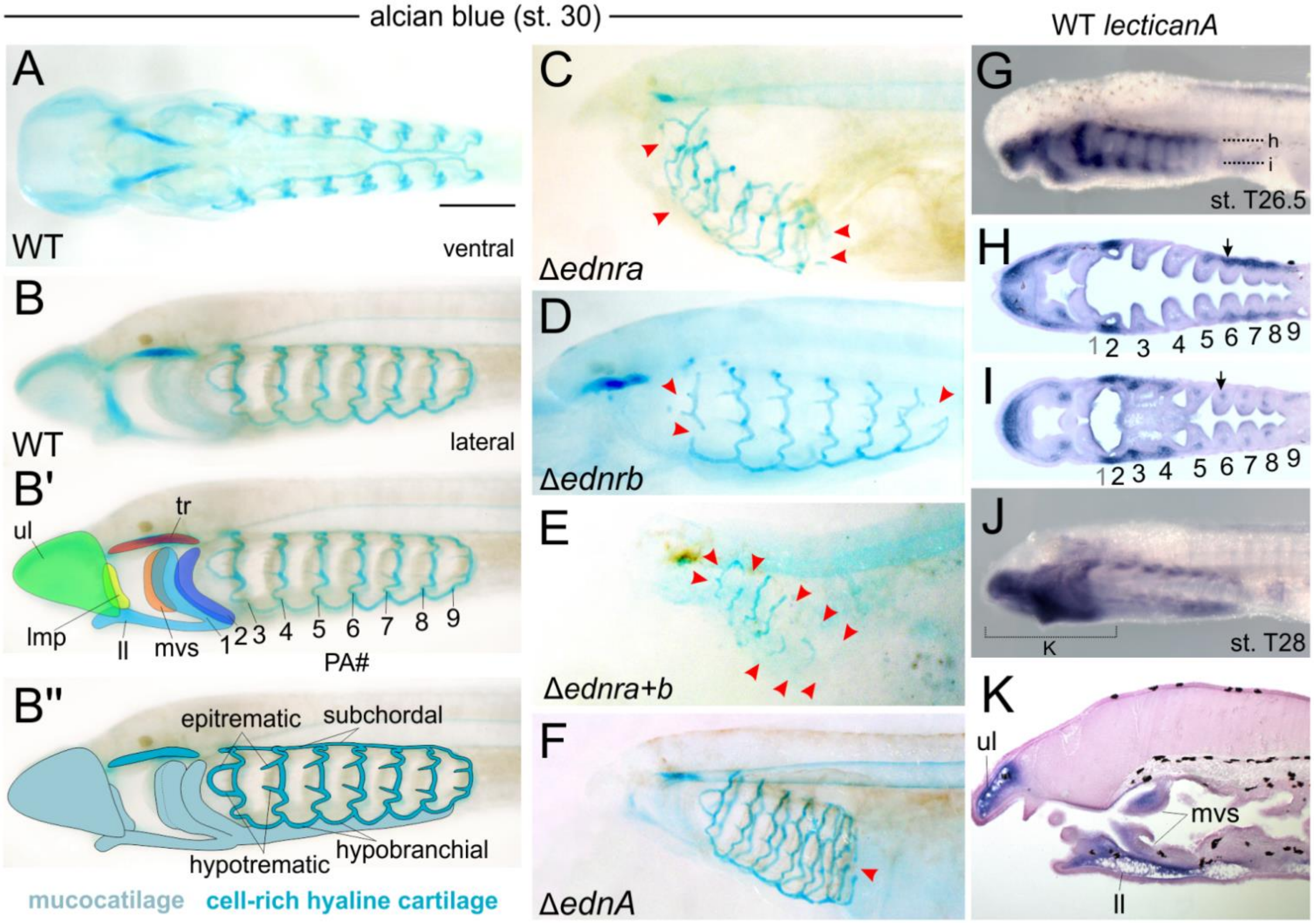


**Supplemental Figure 1 | *Petromyzon marinus* wildtype and mutant larval alcian blue stained head skeletons and *lecticanA* expression.** Anterior to left in all panels. **A**, WT ventral view at stage T30. **B**, WT lateral view of the same specimen, with skeletal elements and cartilage types labeled in b' and b'', respectively. in b', regions of the oral skeleton are delineated. In b'', "epitrematic" indicates the epitrematic processes on PAs 3 and 4, though these exist on all branchial arches. "Hypotrematic" indicates the hypotrematic processes of PAs 3 and 4, though these exist on all branchial arches. "hypobranchial" indicates the hypobrancial cartilage connecting PAs 4, 5, and 6, but exists between all branchial arches. "Subchordal" indicates the subchordal cartilage connecting PAs 4, 5, and 6, but exists between all branchial arches. **C-F**, Δ*ednra,* Δ*ednrb*, Δ*ednra+b*, Δ*ednA* head skeleton phenotypes at stage T30. Red arrowheads highlight some regions where cartilages are missing or separated. **G-K**, *lecticanA* expression summary in *P. marinus*. This gene is homologous to gnathostome lecticans (such as *aggrecan*) and like those genes it is expressed in neural crest-derived mesenchyme before (e.g. arrows in G and H) and during chondrogenesis (e.g. expression in J). ll, lower lip; lmp, lateral mouth plate; mvs, medial velar skeleton; PA#, pharyngeal arch (numbered individually); tr, trabecular; ul, upper lip.

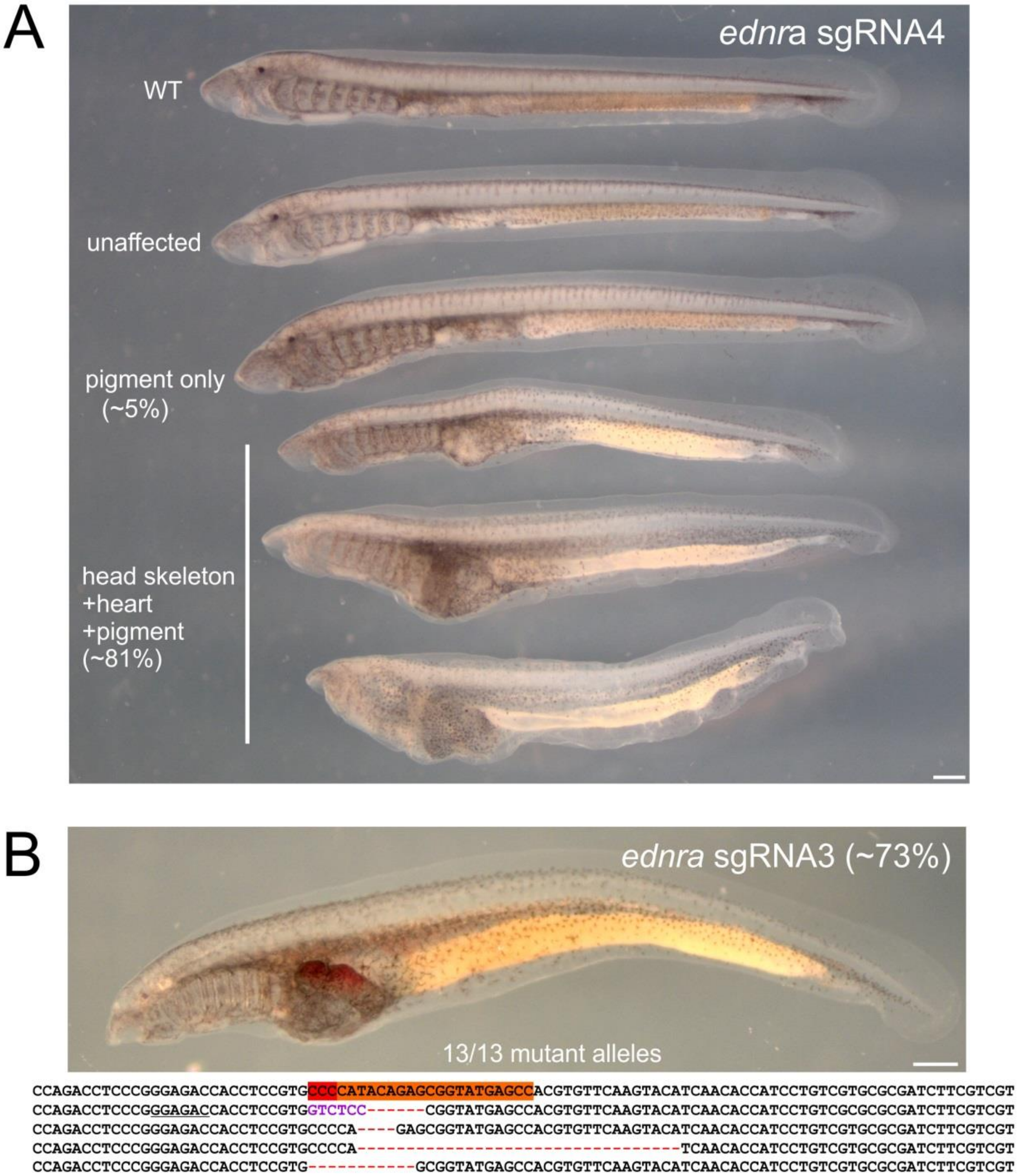


**Supplemental Figure 2 | *Petromyzon marinus* Δ*ednra* phenotype and genotype summary. A**, *ednra* sgRNA4 phenotype gradient at stage T30, which was observed for both *ednra* sgRNAs. **B**, an example of a genotyped animal from injected with *ednra* sgRNA3 which returned 13/13 indel alleles. Target site is shown in orange with a red PAM. Purple nucleotide string indicates an insertion that reflects endogenous sequence near the lesion on the reverse strand (underlined nucleotide string). An insertion is stacked inside of the lesion on the 5' end. Both scale bars represent 500 μm.

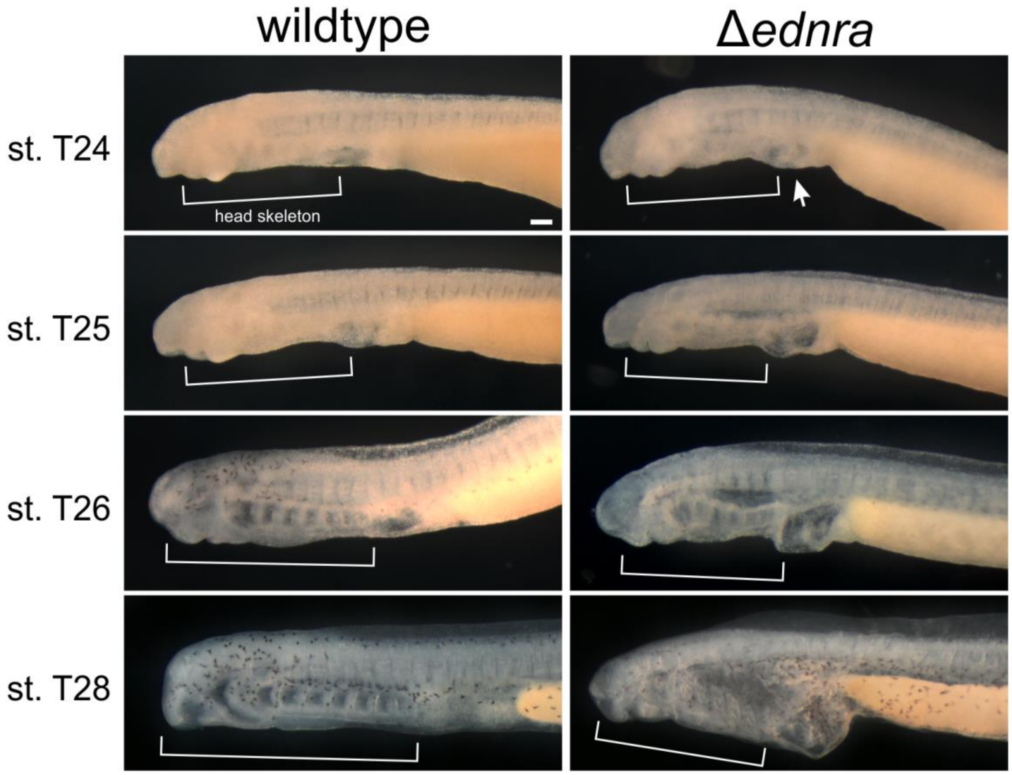


**Supplemental Figure 3 | *Petromyzon marinus* Δ*ednra* phenotype progression.** A staging series of Δ*ednra* illustrating the typical manifestation of the severe phenotype. At stage T24, a slight heart edema is usually apparent (arrow). From this point, the reduction in head skeleton size becomes progressively more dramatic through time relative to WT (brackets mark the anterior and posterior boundaries of the skeletogenic mesenchyme). Scale bar represents 100 µm and applies to all images.

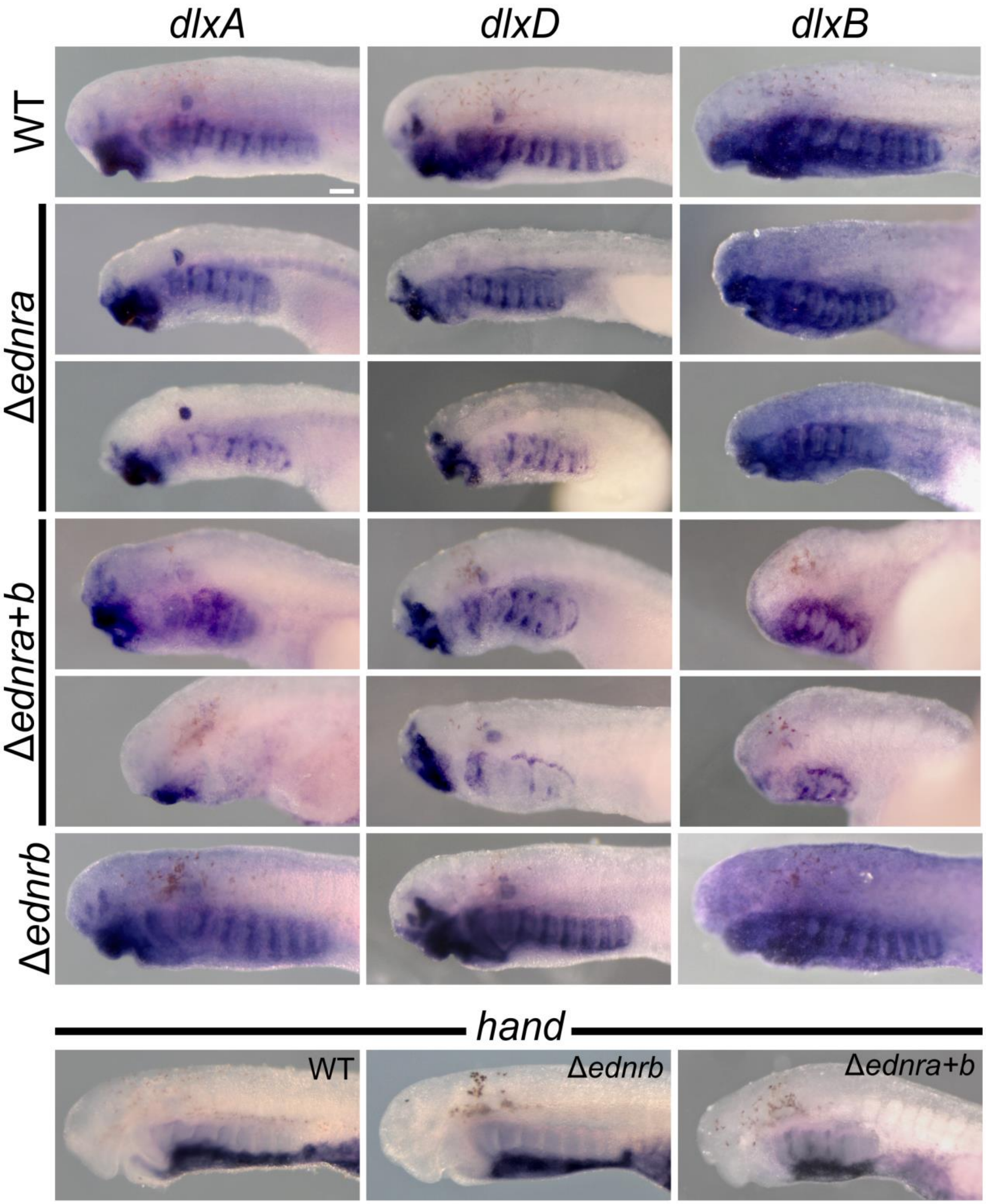


**Supplemental Figure 4 | *Petromyzon marinus dlxA, -D, -B,* and *hand* expression in Δ*ednr* lampreys at stage T26.5.** Left lateral views with anterior to left in all panels. All panels are to scale with each other. Treatment and gene assayed are indicated in the figure. Scale bar represents 100 µm and applies to all images.

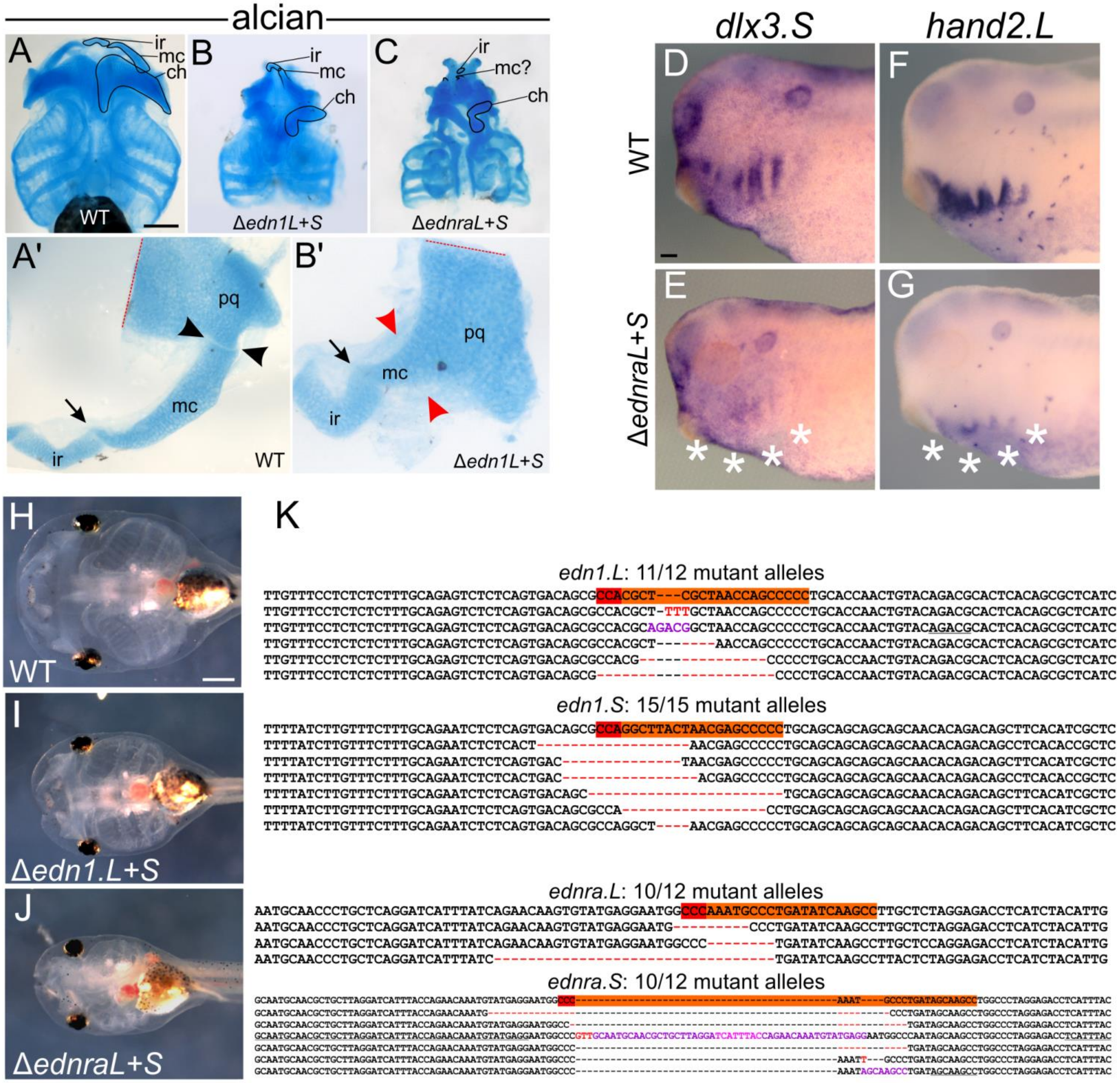


**Supplemental Figure 5 | *Xenopus laevis* Δ*ednra* and Δ*edn1* head skeleton defects and genotyping. A-C**, Δ*edn1L+S* (B) and Δ*ednraL+S* (C) show hypomorphic head skeleton elements relative to WT (A) at stage 48. Δ*edn1L+S* is typically less severe, with a discernible Meckel's cartilage (labeled "mc") present but fused to the palatoquadrate ("pq"), thus lacking a primary jaw joint (black arrowheads in a', red arrowheads in b'), similar to zebrafish *sucker* (*edn1*) mutants. However in Δ*ednraL+S,* Meckel's cartilage is highly reduced, and frequently unrecognizable, only visible as a bump on the palatoquadrate in most cases. The infrarostral ("ir") is always detectable, and no fusion of this element to Meckel's cartilage was ever observed (i.e. the intramandibular joint appears unaffected by a loss of *edn1* or *ednra*, arrows in a' and b'). The ceratohyal ("ch") was highly reduced in both perturbations. The branchial arch skeleton (comprising pharyngeal arches 3-6), though slightly reduced, kept its overall structure more in-tact than PA1 and PA2 derivatives (e.g. Meckel's and the ceratohyal). Ventral views with anterior to top in a-c, dissected views in a' and b', red dotted line in a' and b' indicates a cut made during dissection through the palatoquadrate. Scale bar in a represents 1 mm and applies to b and c. a' and b' are not to scale with each other. **D-G**, *dlx3.S* and *hand2.L* expression are reduced in Δ*ednraL+S* (asterisks) at stage 33. Lateral views with anterior to left in d-g. Scale bar in d represents 100 µm and applies to e-g. **H-K**, genotyping examples of Δ*edn1L+S* (I, K) and Δ*ednraL+S* (J, K) larvae. The alleles shown in k are derived from the animals pictured in i and j. Target sites are shown in orange with a red PAM. h-j show ventral views with anterior to the left. Pink and purple nucleotide strings indicate inserted sequences that are also observed at the endogenous locus near the lesion on the forward strand (underlined nucleotide strings). Red nucleotide strings represent insertions without an obvious source. Insertions are stacked inside of each the lesions on the 5' end.

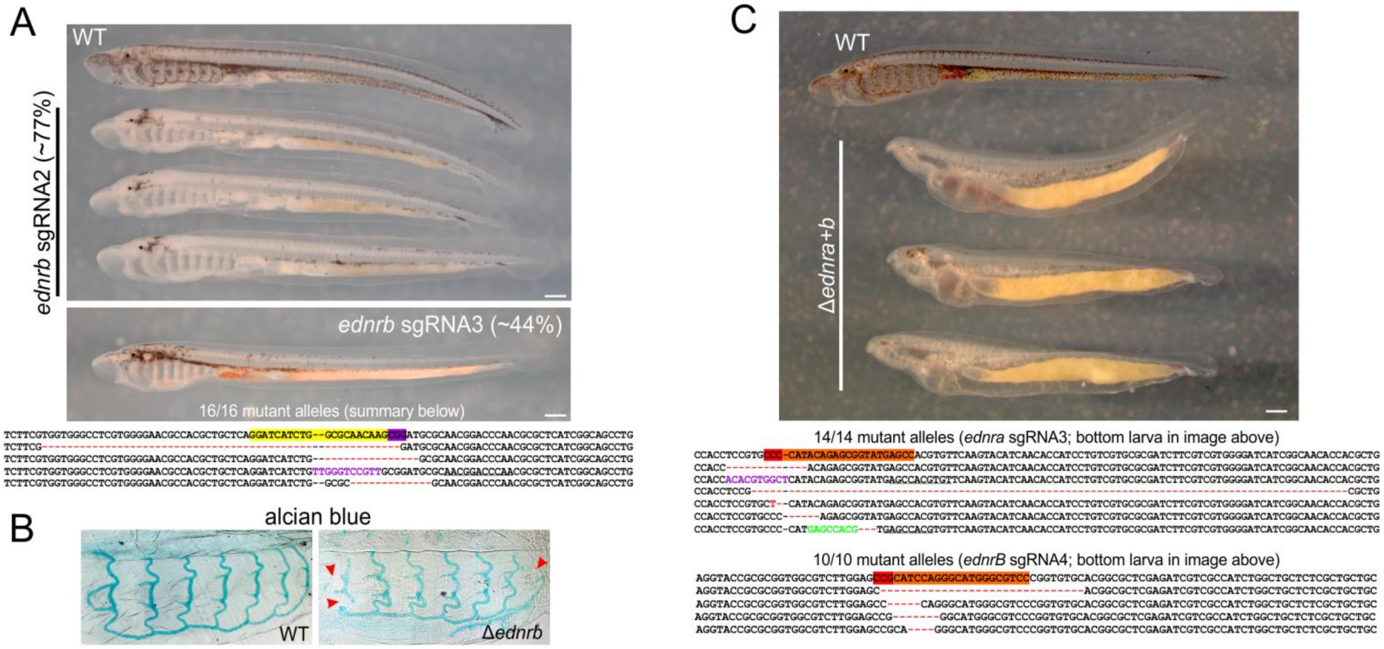


**Supplemental Figure 6 | *P. marinus* Δ*ednrb* and Δ*ednra+b* phenotypes and genotyping. A**, Left lateral images of affected *ednrb* mutants at stage T30. 100% mutant alleles were returned for the indicated individual. Target site for sgRNA3 is shown in yellow with a purple PAM. Four example alleles are shown. An insertion from the reverse strand is shown in purple, its ‘source’ is underlined. The insertion is stacked inside of the lesion on the 5’ end. Both scale bars represent 500 µm. **B**, alcian blue staining reveals slight skeletal disruptions in Δ*ednrb* at stage T30 (red arrowheads)*.* **C**, examples of Δ*ednra+b* at stage T30 that were genotyped and all found to harbor a majority of mutant alleles. A summary of the alleles found in the third individual are shown. Target sites are shown in orange with a red PAM. Purple (forward) and green (reverse) nucleotide strings indicate insertions of sequences that are also observed at each endogenous locus near the lesion (underlined nucleotide strings). Red nucleotide strings represent insertions without an obvious source. Insertions are stacked inside of each the lesions on the 5’ end. Scale bar in c represents 500 µm.

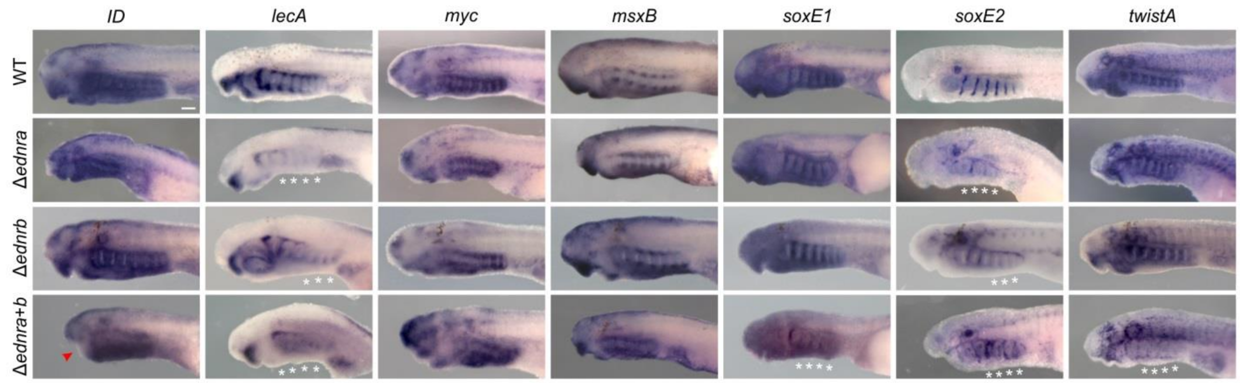


**Supplemental Figure 7 | *P. marinus* Δ*ednr myc, msxB, soxE1, twistA, and ID* gene expression at T26.5.** Lateral views with anterior to left in all panels. Treatment and gene assayed are indicated in the figure. Overall, Δ*ednra* and Δ*ednrb* are most consistent in their gene expression patterns, despite all domains being shrunken in Δ*ednra*. Converseley, Δ*ednra+b* displays some disruptions in gene expression not observed in either single receptor perturbation (red arrowheads indicate specific regions of recused expression, white stars indicate large scale loss of gene expression).

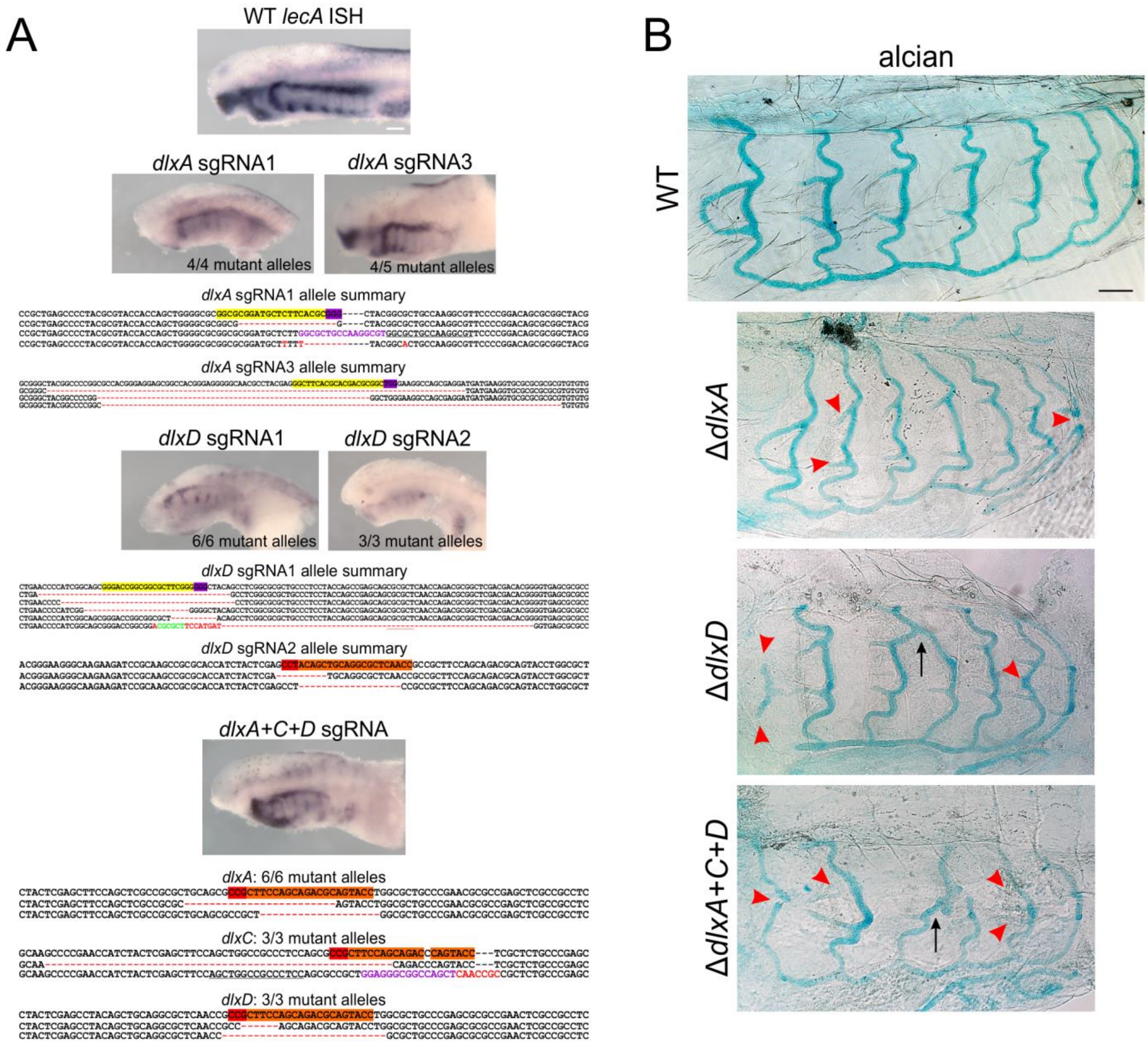


**Supplemental Figure 8 | *Petromyzon marinus* Δ*dlx* genotyping post-ISH and alcian blue staining. A**, examples of post-ISH genotyping for each *dlx* locus targeted for mutagenesis at stage T26.5. Gene, target site, and measured frequency of indel alleles are all indicated in the figure for each individual. Forward strand target sites are shown in yellow with a purple PAM, reverse strand target sites are shown in orange with a red PAM. Purple and green nucleotide strings represent insertions that reflect endogenous sequence in the forward and reverse orientation, respectively, near the lesion (underlined nucleotide strings). Red nucleotide strings represent insertions without an obvious source. Insertions are stacked inside of each the lesions on the 5' end. All ISH images are to scale with each other. **B**, alcian blue staining at stage T30 reveals truncations and gaps in the intermediate head skeleton (arrowheads), and occasional fusions of branchial arches (arrows) in Δ*dlx* lampreys. Scale bar in b represents 100 µm.

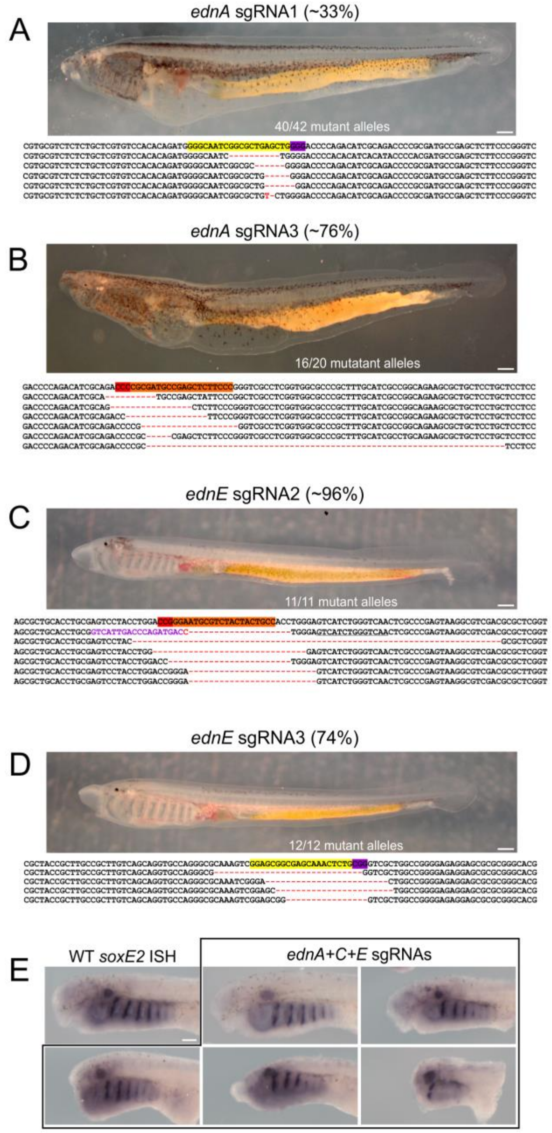


**Supplemental Figure 9 | *Petromyzon marinus* Δ*ednA* and Δ*ednE* phenotype and genotype summary. A-D,** examples of genotyped material and the loci returned for each specimen. Gene, target site, and measured frequency of indel alleles are all indicated in the figure for each individual. Forward strand target sites are shown in yellow with a purple PAM, reverse strand target sites are shown in orange with a red PAM. Purple nucleotide string indicates an insertion that is observed at the endogenous locus in the forward orientation near the lesion (underlined nucleotide string). Red nucleotide strings represent insertions without an obvious source. Insertions are stacked inside of each the lesions on the 5' end. Scale bars represent 500 µm. **E**, *ednA*, -*C*, and -*E* sgRNAs were coinjected, and the material was subjected to *soxE2* ISH at stage T26.5.

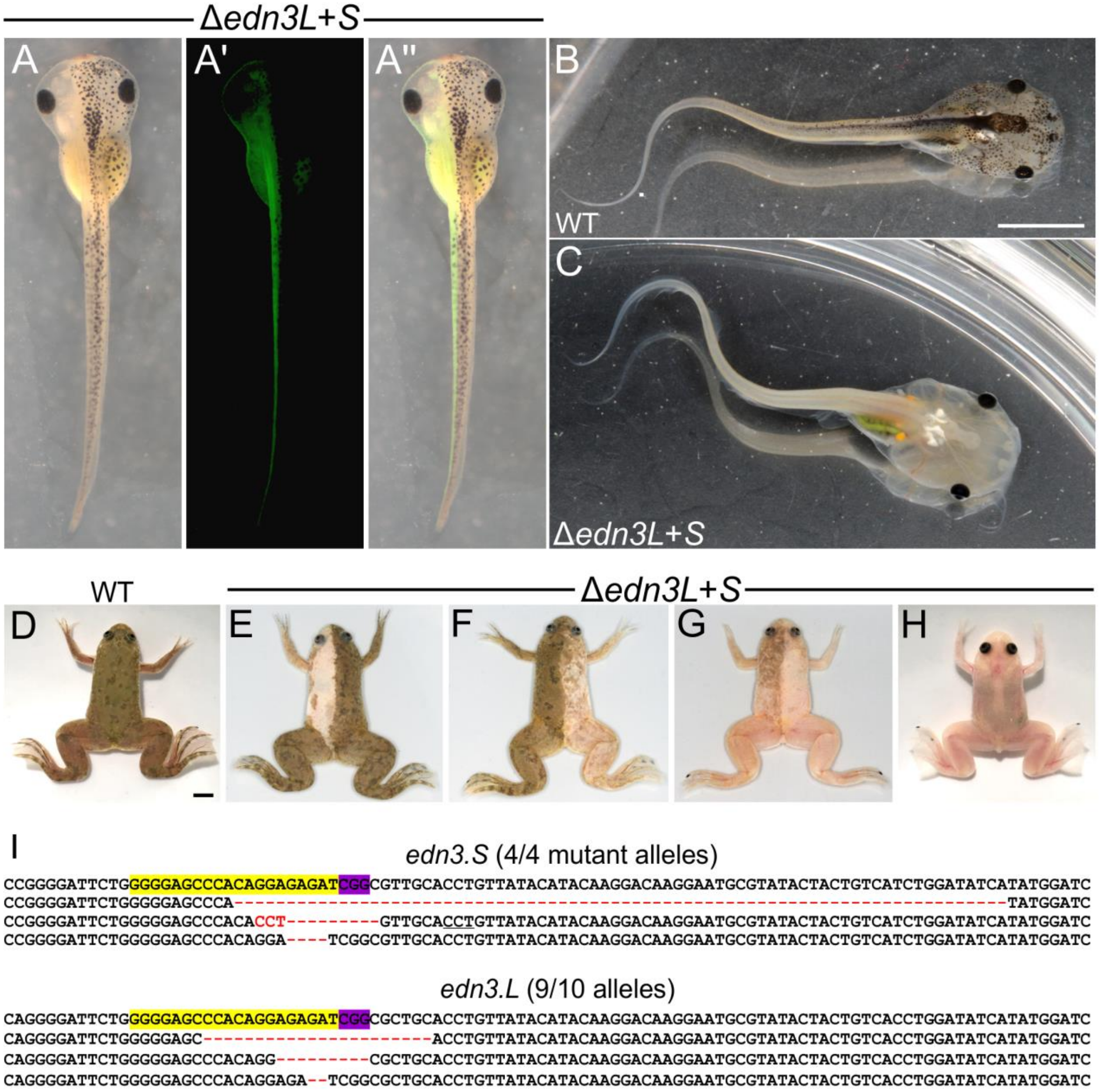


**Supplemental Figure 10 | *Xenopus laevis* Δ*edn3* phenotype and genotype summary. A-H**, Δ*edn3L+S* (A, C, E-H) have reduced neural crest-derived pigment cells (including at least melanophores and iridophores) relative to WT (B, D), or in the case of a, e, f, and g, the uninjected half of the specimen (these animals were injected unilaterally at the 2 blastomere stage, a' shows the lineage tracer GFP [coinjected as mRNA], a'' shows an overlay of a and a'). As expected, pigmentation loss in the eye was never observed (as eye pigmentation is not derived from the neural crest), and the black coloration of the claws always remains, which is also observed in *tyrosinase* mutants (observed by both our lab group and that of Yonglong Chen, personal communication). All images show dorsal views, anterior to top in a, d-h, and to the right in b and c. Scale bar in b represents 5 mm and applies to c. Scale bar in d also represents 5 mm and applies to e-h. **I**, genotyping of a leucistic tadpole revealed a high rate of mutant alleles across both the "Short" and "Long" homoeologs. Target sites are shown in yellow with a purple PAM site.

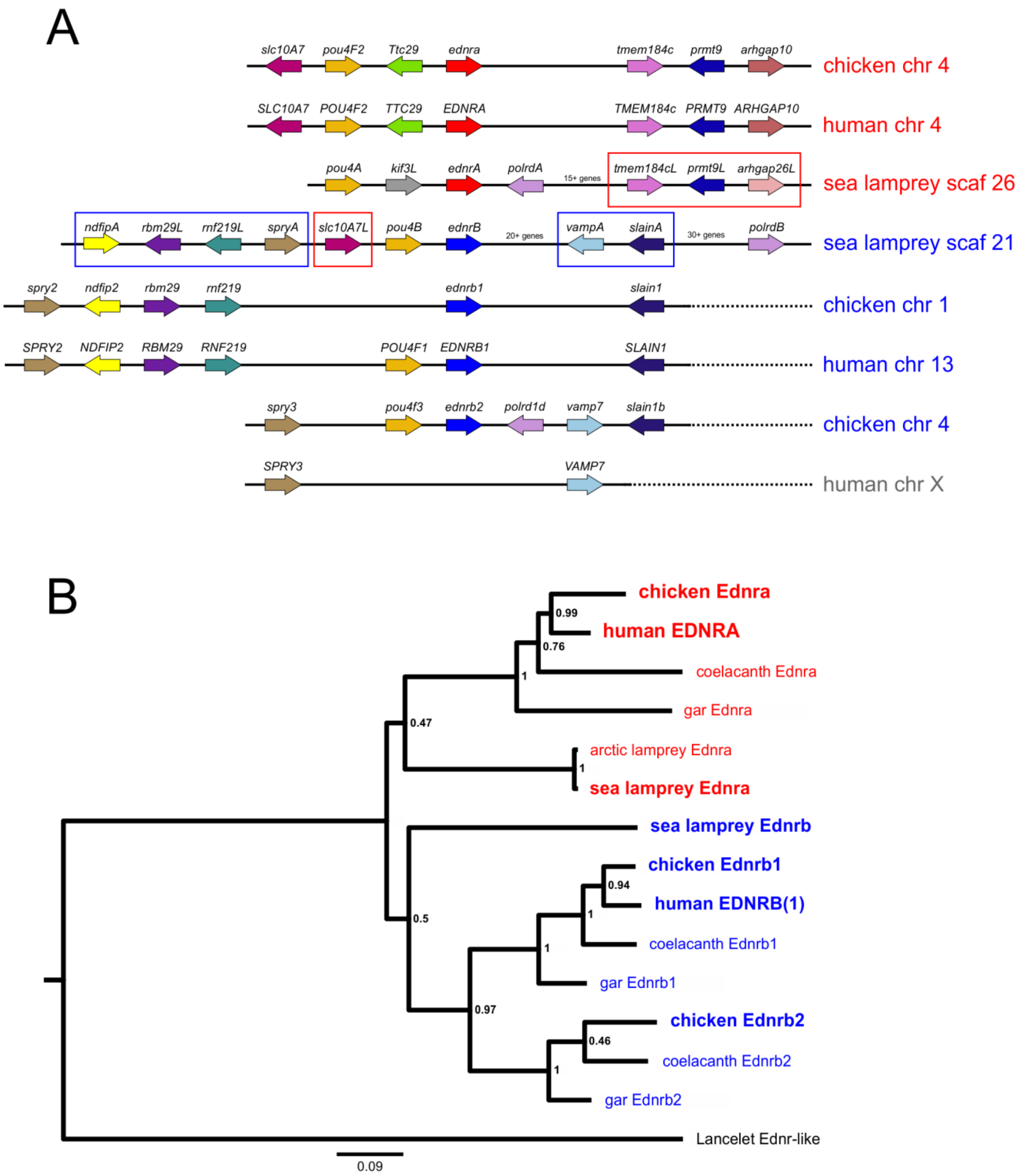


**Supplemental Figure 11 | *ednr* synteny and amino acid sequence comparison. A**, synteny of *ednr*s. All *ednr* loci are shown for chicken (*Gallus gallus*), human (*Homo sapiens*), and sea lamprey (*Petromyzon marinus*), as well as lancelet (*Branchiostome belcheri*) Ednr-like. Red and blue boxes indicate genes or groups of genes that were only observed at the *ednra* or *ednrb* loci, respectively, across species. Chicken and human information is after Braasch and Schartl, 2015. Sea lamprey genomic information is derived from the germline genome published via JJ Smith *et al*., 2018. **B**, amino acid tree made using Maximum Likelihood on a ClustalW alignment, derived from a subset of sequences used in Square et al., 2016. Bootstrap proportions (n=100) are indicated at each node. See Supplemental table 5 for accession numbers.

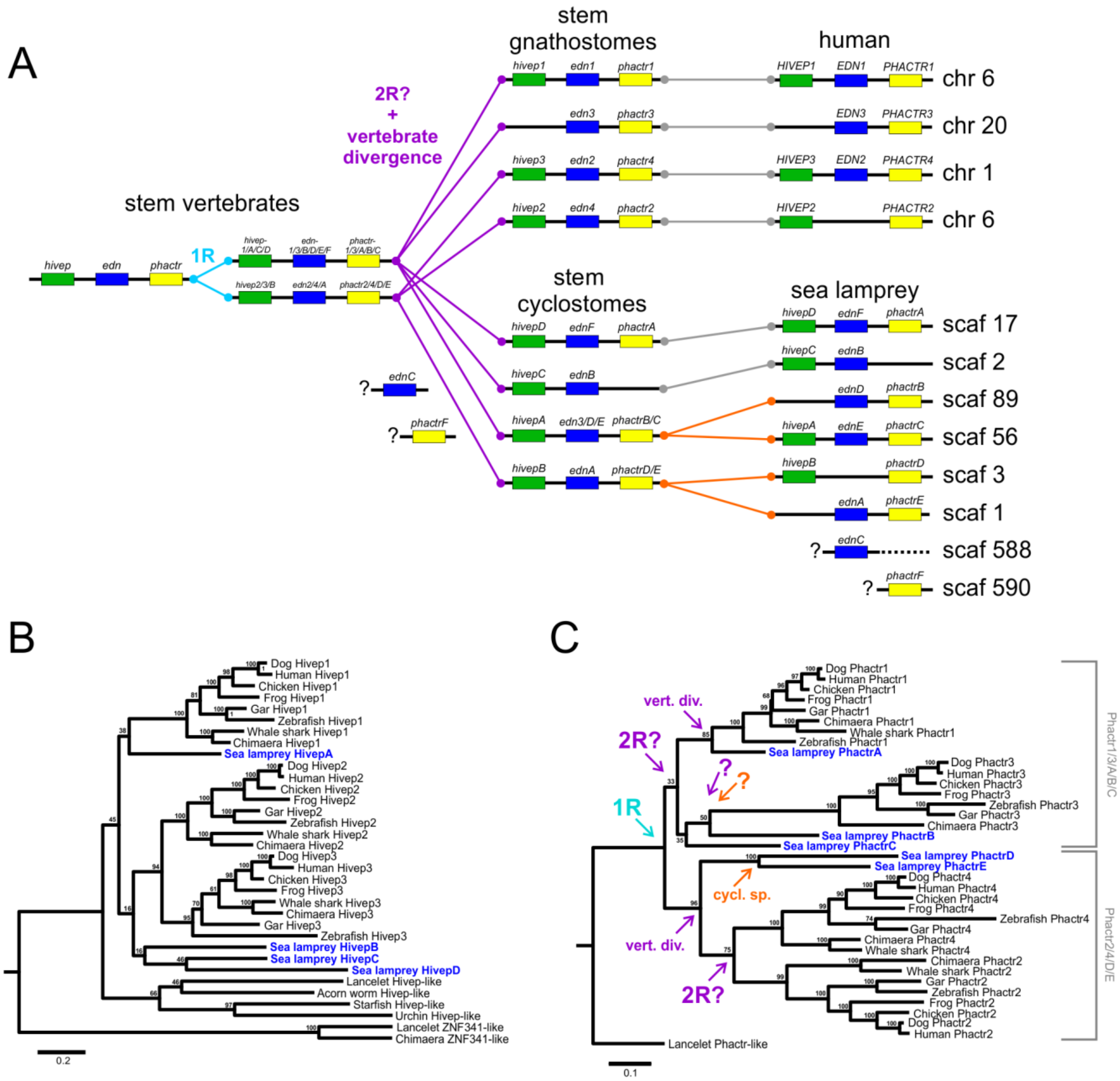


**Supplemental Figure 12 | *edn* relationships. A**, Most vertebrate *edn* genes are located between *hivep* and *phactr* paralogs, which were previously used by Braasch *et al*., 2009 to determine the relationships of jawed vertebrate *edn*s. We found that sea lamprey *edn* genes are also associated with *hivep* and/or *phactr* genes in most cases, and used the sequence similarity of these gene products to address the relatedness of the loci housing the *edn* genes. **B and C**, Hivep and Phactr amino acid comparisons. Bootstrap frequencies are indicated at each node (out of 100 bootstrap replicates). See Supplemental table 5 for sequence information and accession numbers.

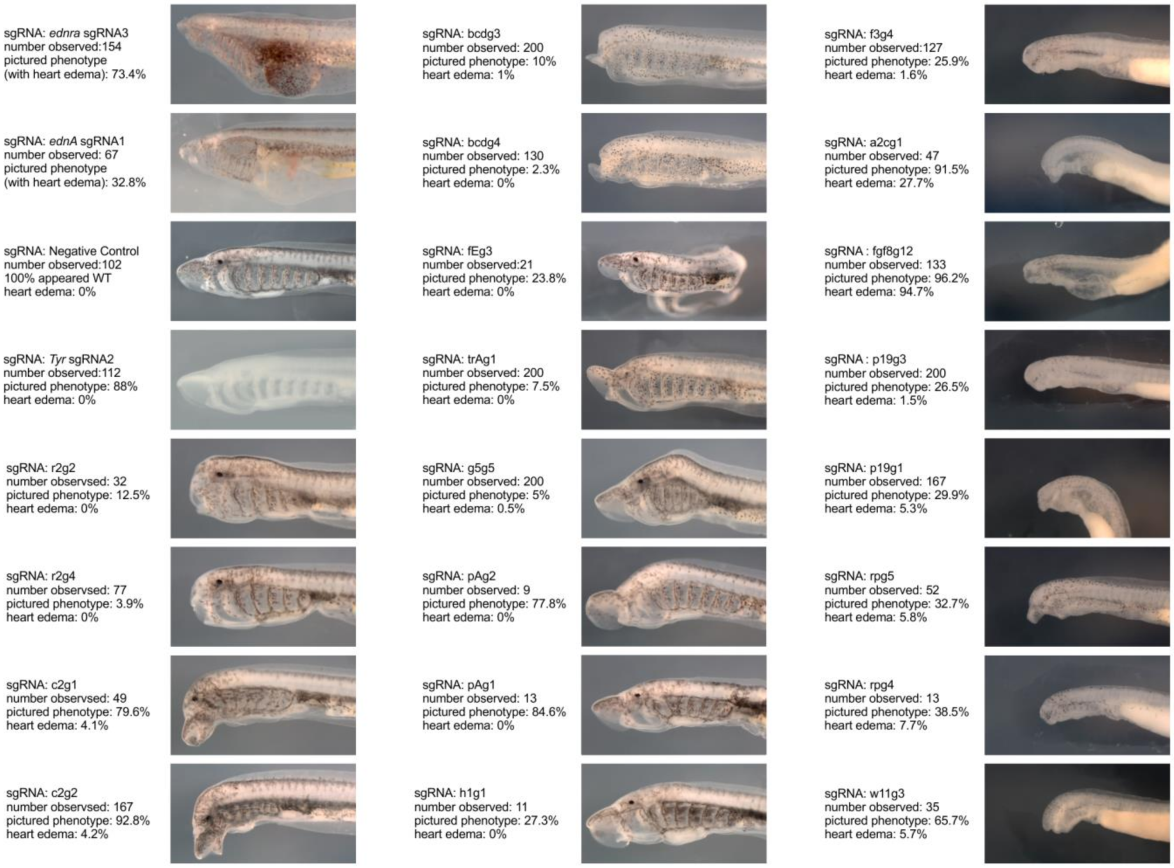


**Supplemental Figure 13 | control injections.** A litany of other developmental regulators was targeted with Cas9 injections. These caused a wide range of phenotypes at T30 and T26.5, though heart edema was rarely detected. This indicates that the phenotypes described by this work are likely specific to the genes of interest, rather than non-specific side effects of sgRNA + Cas9 protein injections and the resulting genomic lesions.
